# Increased chloroplast area in the rice bundle sheath through cell specific perturbation of brassinosteroid signalling

**DOI:** 10.1101/2024.08.14.607565

**Authors:** Lee Cackett, Leonie H. Luginbuehl, Ross-William Hendron, Andrew R. G. Plackett, Susan Stanley, Steven Kelly, Julian M. Hibberd

## Abstract

In the leaves of C_3_ species such as rice, mesophyll cells contain the largest compartment of photosynthetically active chloroplasts. In contrast, plants that use the derived and more efficient C_4_ photosynthetic pathway have a significant chloroplast compartment in both bundle sheath and mesophyll cells. Accordingly, the evolution of C_4_ photosynthesis from the ancestral C_3_ state requires an increase in the chloroplast compartment of the bundle sheath. Here we investigated the potential to increase chloroplast compartment size in rice bundle sheath cells by manipulating brassinosteroid signalling. Treatment with brassinazole, a brassinosteroid biosynthesis inhibitor, increased leaf chlorophyll content and increased the number but decreased the area of chloroplasts in bundle sheath cells. Constitutive overexpression of the *BRASSINAZOLE RESISTANT 1* (*OsBZR1*) transcription factor increased bundle sheath chloroplast area by up to 45% but plants became chlorotic. However, when *OsBZR1* was placed under the control of a bundle sheath specific promoter, the negative effects on growth and viability were removed whilst chloroplast area still increased. In summary, we report a role for brassinosteroids in controlling chloroplast area and number in rice and conclude that cell-specific manipulation of brassinosteroid signalling can be used to manipulate the chloroplast compartment in rice bundle sheath cells.

## Introduction

Increasing crop yield is considered imperative to feed a growing population and improving photosynthetic efficiency is recognised as one possible approach to achieve this (Smith et al., 2023). In land plants photosynthesis takes place in chloroplasts, the development of which is initiated by the perception of light and is modulated by various hormones (Cackett et al., 2022). This interplay allows the chloroplast content of each cell type to be tuned to the needs of the cell. For example, in C_3_ species such as rice (*Oryza sativa*), carbon fixation occurs primarily in mesophyll cells that are densely packed with chloroplasts (Sage & Sage, 2009). Although bundle sheath cells in C_3_ plants also contain chloroplasts, the proportion of cell volume they occupy is much lower than in mesophyll cells (Sage & Sage, 2009). In contrast, C_4_ plants such as maize (*Zea mays*) contain a greatly enhanced chloroplast volume in bundle sheath cells (Lee et al., 2023). This allows photosynthetic reactions to be partitioned between the mesophyll and bundle sheath such that a biochemical pump concentrates CO_2_ in bundle sheath cells where RuBisCO accumulates. This C_4_ cycle reduces oxygenation of RuBisCO and the subsequent photorespiratory reactions, enabling photosynthetic efficiency to be increased by up to 50% (Sage et al., 2012). Understanding the differences in chloroplast biogenesis between cell types is therefore relevant to attempts to engineer C_3_ leaves such that they operate a C_4_-like photosynthesis.

Chloroplast biogenesis is primarily controlled by transcriptional regulators belonging to the *GOLDEN2-LIKE* (*GLK*), *GATA NITRATE-INDUCIBLE CARBONMETABOLISM- INVOLVED* (*GNC*) and *CYTOKININ RESPONSIVE GATA FACTOR 1* (*CGA1*) families (reviewed by Cackett et al., 2022). Recently, an additional regulator from the *RR-TYPE MYOBLASTOMA RELATED* (*RR-MYB*) family has been identified (Frangedakis et al., 2024) but it is not yet known how it responds to signals inducing chloroplast biogenesis, nor whether other transcription factors are involved. Overexpression of *GLK*s in multiple species increases chlorophyll and chloroplast production and can stimulate this in tissues that normally contain a very limited chloroplast compartment (Kobayashi et al., 2012, 2013; Nakamura et al., 2009; Wang et al., 2017). Constitutive overexpression of *ZmG2* in rice grown in the field increased photosynthesis, vegetative biomass, and grain yield (Li et al., 2020). Overexpression of *GNC* and *CGA1* increased chloroplast planar area in Arabidopsis (*Arabidopsis thaliana*) (Hudson et al., 2011) and rice bundle sheath cells (Hudson et al., 2013; Lee et al., 2021; Lim et al., 2024). However, neither overexpression of *GLK* or *CGA1* stimulated chloroplast biogenesis in the rice bundle sheath to the extent that their chloroplast content matched that of C_4_ sorghum or maize. Whilst unknown regulators may control the enhanced biogenesis of bundle sheath chloroplasts in C_4_ species, it is also plausible that known components initiating this process are responsible, but the complete network of control has not yet been elucidated.

The brassinosteroid signalling pathway acts to repress chlorophyll accumulation and chloroplast biogenesis in the dark and is inhibited after light is perceived and de-etiolation is induced. In the dark, brassinosteroids act with PHYTOCRHOME INTERACTING FACTORS (PIFs) to negatively regulate photosynthesis gene expression (Oh et al., 2012). Upon exposure to light, PIFs are degraded in response to phytochrome signalling and the induction of *GNC* expression represses BR signalling, allowing activation of chloroplast biogenesis (reviewed by Cackett et al., 2022). In the dark, Arabidopsis BR-related mutants such as *det2, dwf4*, *cpd, bri1* and *bin2* show characteristics of de-etiolation including differentiated chloroplasts, short hypocotyls, development of true leaves and expression of light-regulated genes (Azpiroz et al., 1998; Chory et al., 1991; Clouse et al., 1996; Li et al., 2001; Szekeres & Né, 1996; Tachibana et al., 2022). The transcription factor BRASSINAZOLE RESISTANT 1 (BZR1) mediates the brassinosteroid-modulated negative control of photomorphogenesis by repressing genes involved in light-signalling and chloroplast development including photoreceptors such as phytochrome B and phototropin1, transcription factors such as *GATA2, GATA4*, *GLK1&2* and photosynthesis genes associated with chlorophyll biosynthesis (Luo et al., 2010; Sun et al., 2010; Wang et al., 2020; Yu et al., 2011). It is thought that the BZR1 mediated repression of chlorophyll biosynthesis avoids overaccumulation of protochlorophyllide in the dark so that when light is perceived photooxidative damage is minimized and greening promoted (Wang et al., 2020).

The role of brassinosteroids and BZR1 during de-etiolation, organ development, cell elongation and chlorophyll accumulation is well documented in Arabidopsis and, to a lesser extent, in rice. However, to our knowledge there are no analyses in rice demonstrating if or how brassinosteroids control chloroplast biogenesis in terms of size and number per cell. We therefore assessed how pharmacological and genetic perturbations to brassinosteroid signalling affect the planar area and number of chloroplasts in the bundle sheath of rice. Our analysis indicated that brassinosteroids alter the number and area of chloroplasts in the rice bundle sheath during de-etiolation and at later stages of development. Constitutive overexpression of *OsBZR1* resulted in larger chloroplasts in the bundle sheath but had adverse effects on plant health and yield. However, when overexpression of *OsBZR1* was driven by a bundle sheath specific promoter, increased bundle sheath cell chloroplast area was maintained whilst the adverse effects on growth mitigated. Overall, these data are consistent with an approach in which cell-specific manipulation of brassinosteroid signalling could be used to manipulate chloroplast number and size in rice.

## Results

### Brassinosteroids modulate chloroplast size and number in the rice bundle sheath

To initiate an understanding of the role of brassinosteroids in modulating chloroplast biogenesis in rice we applied the active brassinosteroid, brassinolide (BL), or the biosynthesis inhibitor, brassinazole (Brz) to seedlings during de-etiolation. In control plants, greening and expansion of the first leaf were evident after exposure to light as expected (**Fig. 1A**). Consistent with previous analyses (Hong et al., 2002; Mori et al., 2002), BL treatment inhibited de-etiolation such that rice seedlings showed reduced leaf expansion and greening (**Fig. 1A**). Quantification of whole seedling chlorophyll levels confirmed that its accumulation was reduced compared with controls (**Fig. 1B**). In contrast to the BL treatment, seedlings treated with Brz showed an increase in accumulation of chlorophyll after exposure to light compared with controls. Chloroplast content in the bundle sheath cells of the first leaf of controls and treated seedlings were imaged using confocal laser scanning microscopy at 0, 4 and 12 hours after light exposure (**Fig. 1C**). Brz treatment resulted in more chloroplasts per bundle sheath cell compared with controls after 12 hours of light, whereas BL treatment produced no significant difference (**Fig. 1D & E**). Both BL and Brz treatments resulted in smaller bundle sheath chloroplasts compared to control plants, in terms of mean planar area (**Fig. 1F & G**). Bundle sheath cell size decreased compared to the control as a result of the BL treatment and was unchanged in the Brz treatment compared with the control (**Supp. fig. 1**). Overall, these data are consistent with previous studies from other species reporting that brassinosteroids modulate chlorophyll accumulation during de-etiolation but also indicate that brassinosteroids can control chloroplast number and planar area.

**Figure 1:**
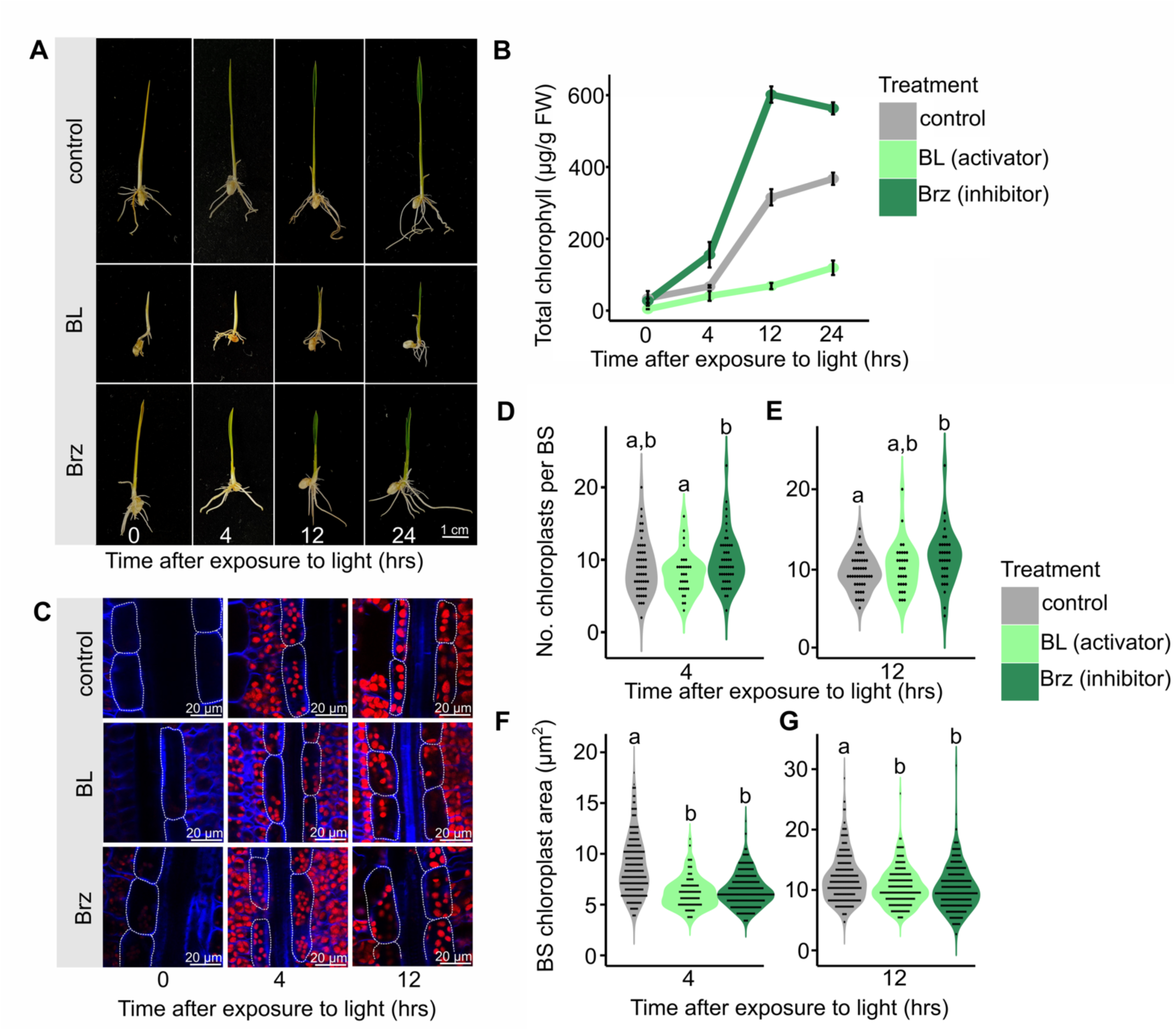
Brassinosteroids modulate chlorophyll accumulation and bundle sheath chloroplast area and number during de-etiolation. Seeds were germinated in water and transferred in the dark to ½ MS-agar media with or without 10 µM brassinolide (BL) or 10 µM brassinazole (Brz). After 4 days seedlings were transferred to light and shoot tissue harvested 0, 4, 12 and 24 hours later for chlorophyll quantification and imaging using confocal laser scanning microscopy. **A:** Representative images of control and BL/Brz treated seedlings during de-etiolation. **B:** Mean chlorophyll content during de-etiolation. Data are from 5 biological repeats for each timepoint in each treatment. **C:** Confocal images of bundle sheath cells and chloroplasts during de-etiolation. Red and blue channels indicate chloroplasts and cell walls respectively. White dotted lines highlight bundle sheath cells. **D** and **E:** Number of chloroplasts per bundle sheath cell (BS) in control and BL/Brz treated seedlings 4 (D) and 12 (E) hours after exposure to light. Data are derived from confocal microscopy and from at least thirty bundle sheath cells for each timepoint in each treatment. **F** and **G:** Bundle sheath cell chloroplast area in control and BL/Brz treated seedlings 4 (F) and 12 (G) hours after exposure to light. Data are derived from confocal microscopy and from at least 150 chloroplasts for each timepoint in each treatment. Letters above violins represent statistically significant differences (*p* ≤ 0.05) in mean values as determined by Fisher LSD post-hoc analysis following a one-way ANOVA.

To determine whether changes to chloroplast number and size were maintained later in development, plants were grown in BL or Brz and leaf 4 harvested once it was fully expanded. Both treatments impacted overall plant growth and development. For example, plants treated with BL developed the same number of leaves as controls, but leaf length was reduced (**Fig. 2A**). Brz caused faster development such that more leaves were evident (**Fig. 2A**), and they contained more veins than controls (**Supp. fig. 2**). Consistent with the initial de-etiolation experiments, chlorophyll content of leaf 4 was reduced by addition of BL (**Fig. 2B**) whilst it had no significant impact on chloroplast area and number in the bundle sheath at this developmental stage (**Fig. 2C, D & E**). Conversely, Brz treatment increased bundle sheath chloroplast number and decreased chloroplast area compared with controls (**Fig. 2C, D & E**) but with no change in chlorophyll content (**Fig. 2B**). Overall, these results indicate that brassinazole, an inhibitor of brassinosteroid biosynthesis, modulates the area and number of chloroplasts in rice bundle sheath cells both during de-etiolation and at later stages of leaf development.

**Figure 2:**
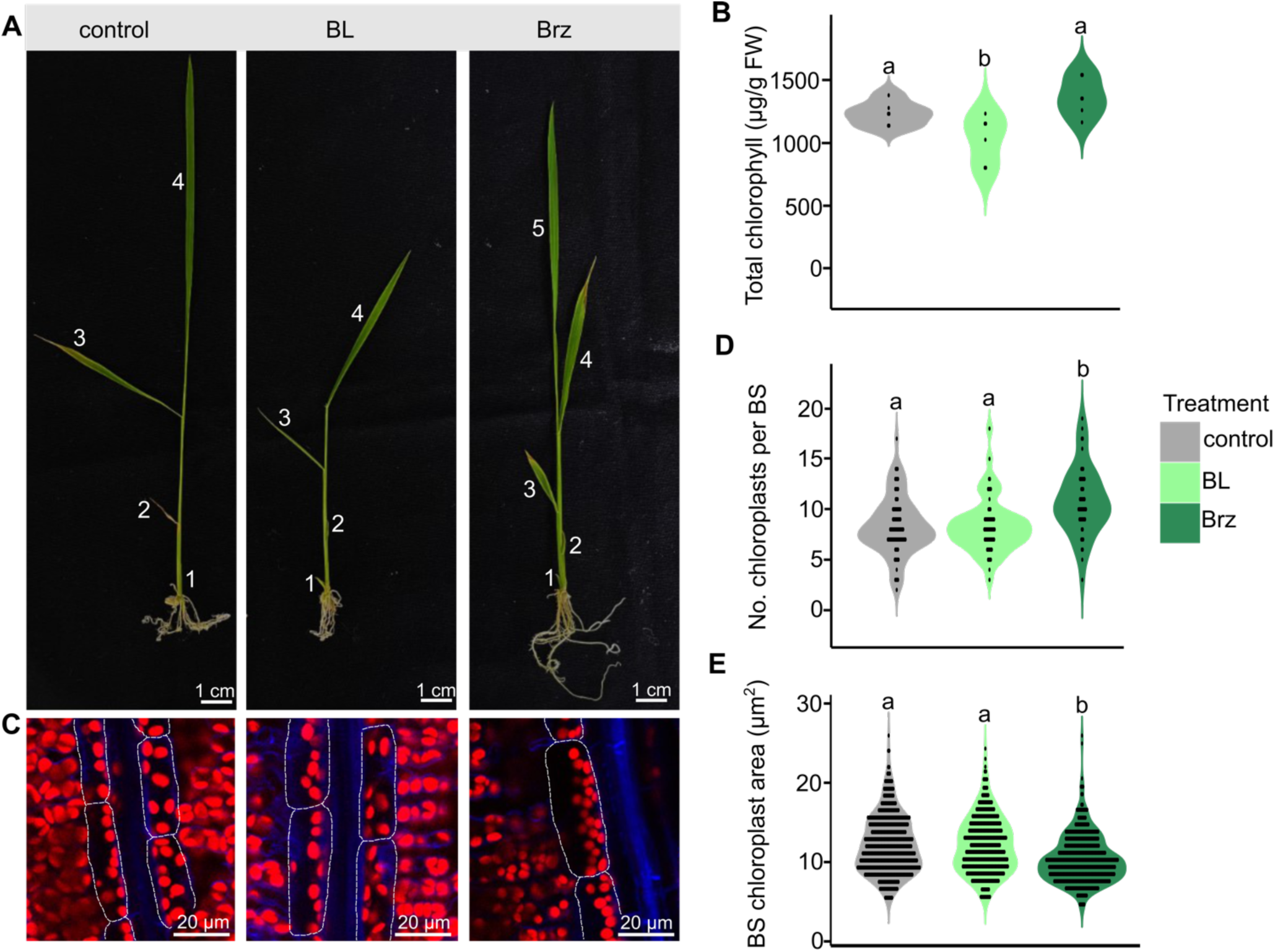
Brassinosteroids modulate chlorophyll accumulation and bundle sheath chloroplast area and number in mature leaves. Seeds were germinated in water and transferred to ½ MS-agar media with or without 10 µM brassinolide (BL) or 10 µM brassinazole (Brz). Leaf 4, once fully expanded, was harvested for chlorophyll quantification and imaging using confocal laser scanning microscopy. **A:** Representative images of control and BL/Brz treated plants. White numbers indicate leaf number in order of appearance. **B:** Chlorophyll content of leaf 4 from control and BL/Brz treated plants, data are from 6 biological repeats for each treatment. **C:** Confocal images of bundle sheath cells and chloroplasts in leaf 4 of controls and BL/Brz treated plants. Red and blue channels indicate chloroplasts and cell walls respectively. White dotted lines highlight bundle sheath cells. **D:** Number of chloroplasts per bundle sheath cell (BS) in leaf 4 from controls and BL/Brz treated plants. Data are derived from confocal microscopy and from at least 70 bundle sheath cells for each treatment. **E:** Bundle sheath cell chloroplast area in leaf 4 from controls and BL/Brz treated plants. Data are derived from confocal microscopy and from at least 550 chloroplasts for each timepoint in each treatment. Letters above violins represent statistically significant differences (*p* ≤ 0.05) in mean values as determined by Fisher LSD post-hoc analysis following one-way ANOVA.

### Constitutive overexpression of OsBZR1 increases chloroplast area in bundle sheath cells but has adverse effects on plant growth

Given that perturbation of BR signalling through exogenous treatments gave rise to changes in chloroplast area and number in rice bundle sheath cells, we sought to determine whether analogous changes could be achieved through genetic activation of BR-responsive gene expression. As BZR1 is the primary transcription factor that mediates BR-responsive gene expression in Arabidopsis (Luo et al., 2010; Sun et al., 2010; Wang et al., 2020; Yu et al., 2011), we chose to investigate whether manipulation of the expression of the orthologous regulatory gene in rice could achieve the desired changes in chloroplast development. Phylogenetic interrogation of the BZR1 gene family revealed that there is a single gene in rice (LOC_ Os07g39220) that is orthologous (i.e. equally related) to BZR1, BES1 (BZR2), BEH1 and BEH2 in Arabidopsis (**Supp. fig. 3**), with no other rice gene homologs in this same clade. Accordingly, we hypothesised that this single gene is likely the primary transcription factor that mediates BR-responsive gene expression in rice. Moreover, previous investigations had implicated this gene in brassinosteroid signalling in rice leading to the naming of the gene *OsBZR1* (Bai et al., 2007). Thus, both overexpression and RNAi lines were generated to alter the expression of Os*BZR1* in the rice leaf.

**Figure 3:**
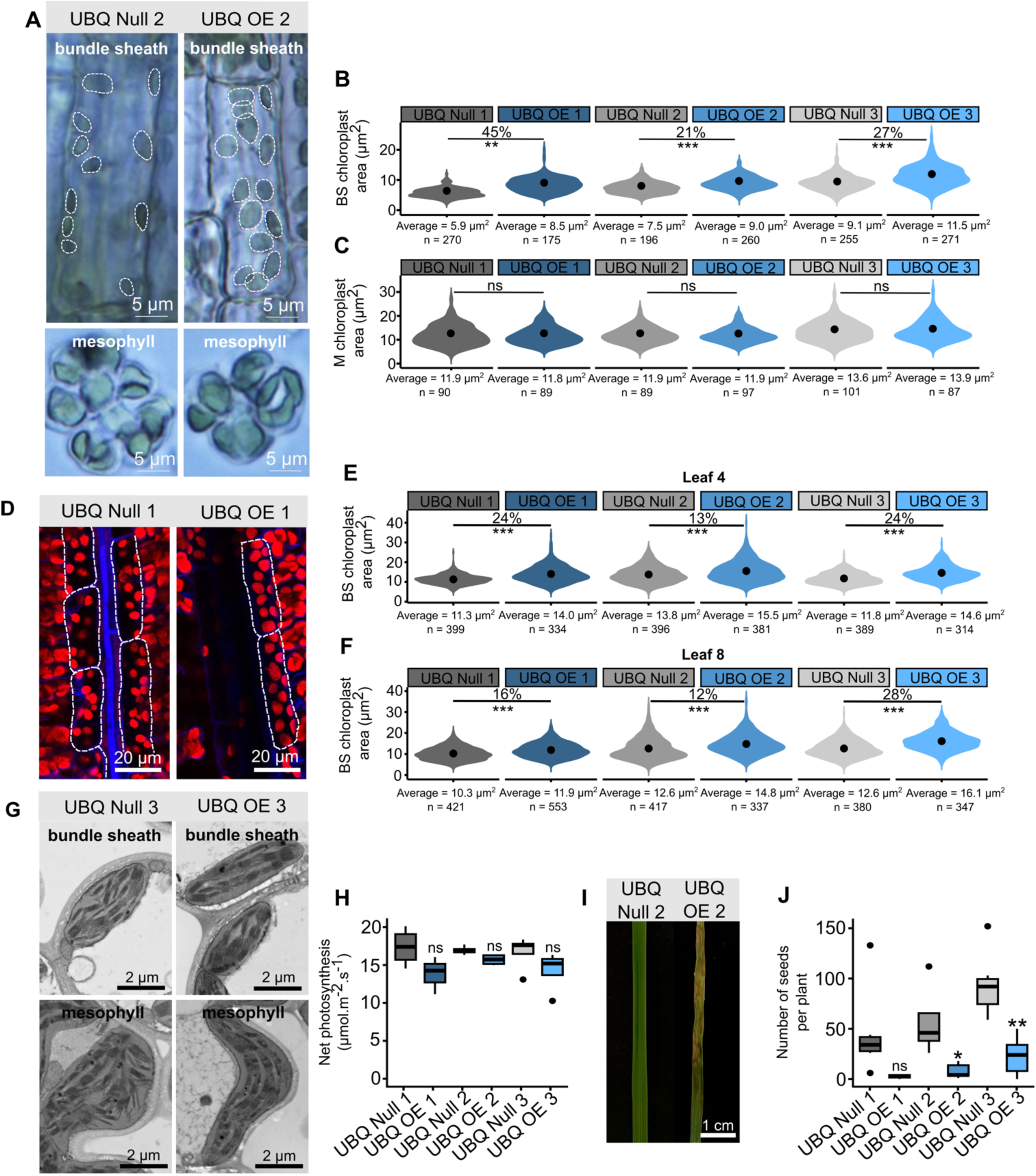
Constitutive overexpression of *OsBZR1* increases the area of chloroplasts in rice bundle sheath cells but impacts plant health. The rice codon optimised sequence for *BZR1* (rco*OsBZR1*) was cloned upstream of the maize *UBIQUITIN* promoter (p*ZmUBI*) and transformed into rice to generate constitutive overexpression lines (UBQ OE). **A:** Representative images from brightfield microscopy of individual bundle sheath and mesophyll cells from leaf 8. White dotted lines highlight individual chloroplasts within each bundle sheath cell. **B and C:** Chloroplast area in bundle sheath (BS) (**B**) and mesophyll (M) (**C**) cells from leaf 8. Chloroplast areas are from quantification of the brightfield microscopy. Percentages above violins indicate the change in chloroplast area of overexpressor lines compared with corresponding nulls. No statistically significant change in chloroplast area is represented by “ns”. The value below each violin is the mean chloroplast area calculated for that line and n represents the number of chloroplasts quantified. Four biological replicates were used for each line. **D:** Images derived from confocal laser scanning microscopy of bundle sheath cells and chloroplasts in leaf 4. Red and blue channels indicate chloroplasts and cell walls respectively. White dotted lines highlight bundle sheath cells. **E and F:** Bundle sheath cell chloroplast area in leaf 4 (**E**) and leaf 8 (**F**). Chloroplasts quantified are from confocal microscopy. Percentages above violins indicate the change in chloroplast area of overexpressor lines compared with corresponding nulls. No statistically significant change in chloroplast area is represented by “ns”. The value below each violin is the mean chloroplast area calculated for that line and n represents the number of chloroplasts assessed. Four biological replicates were used for each line. **G:** Scanning electron microscope (SEM) images of bundle sheath and mesophyll cell chloroplasts from leaf 8. **H:** Rate of net photosynthesis under conditions of growth for overexpressor lines compared with corresponding nulls. Data are from 4 biological replicates for each line. **I:** Representative images of leaf 8 from null and overexpressor plants of the same age depicting increased senescence. **J:** Number of seeds produced by each line. Data are from 4 biological replicates for each line. For figures B, C, E, F, H and J, stars above violins or boxes indicate a statistically significant difference between overexpressor lines compared with corresponding null as determined by independent t-test, where *p* ≤ 0.05 is flagged with one star (*), *p* ≤ 0.01 is flagged with 2 stars (**) and *p* ≤ 0.001 is flagged with three stars (***). No statistically significant change is represented by “ns”.

No reduction in transcript abundance was detected in T_2_ *OsBZR1* RNAi plants (**Supp. fig. 4B**), and so no phenotyping was performed. However, constitutive overexpression using the maize UBIQUITIN promoter was successful. Here, three independent homozygous single copy transgenic lines along with their respective null segregants (lines which had been through the transformation process but do not contain the genetic modification themselves) were identified (**Supp. fig. 5A & B**). RT-qPCR on T_2_ plants was conducted to confirm the level of transgene expression (**Supp. fig. 5C**). Hereafter these lines are referred to as UBQ Null 1, UBQ OE 1, UBQ Null 2, UBQ OE 2, UBQ Null 3 and UBQ OE 3. Brightfield microscopy was used to image isolated bundle sheath and mesophyll cells from fully expanded leaf 8 (**Fig. 3A**). The individual chloroplast planar area was significantly larger for all three overexpression lines compared with the respective nulls in bundle sheath cells (**Fig. 3B**). UBQ OE 1 showed the largest effect with individual chloroplast area being increased by 45% (**Fig. 3B**). The extent to which chloroplast area increased correlated with the degree of *OsBZR1* overexpression within each line (**Supp. fig. 5C**). There were no statistically significant differences in chloroplast area in mesophyll cells in any UBQ OE line (**Fig. 3C**). To confirm these findings with a higher throughput approach (Billakurthi & Hibberd, 2023) allowing a larger number of chloroplasts to be assessed, we next used confocal laser scanning microscopy (**Fig. 3D**). Consistent with the analysis of single cells after brightfield microscopy, this showed that overexpression of *OsBZR1* increased individual chloroplast area in bundle sheath cells in leaf 4 and leaf 8 compared with corresponding null lines (**Fig. 3E and F**). Scanning electron microscopy detected no evident changes in bundle sheath chloroplast ultrastructure between null and overexpression lines (**Fig. 3G**). There were no consistent differences in whole leaf chlorophyll content (**Supp. fig. 6**) and only one of the three UBQ OE lines showed a significant difference in bundle sheath cell size compared with the corresponding null line (**Supp. fig. 7**). The rate of net photosynthesis in young fully expanded leaves was not affected by constitutive overexpression of *OsBZR1* (**Fig. 3H**). Notably, the UBQ OE leaves senesced rapidly soon after maturation (**Fig. 3I**) and the number of seeds produced by all UBQ OE plants was significantly lower than corresponding nulls (**Fig. 3J**). We therefore sought to test whether a more targeted mis-expression of *OsBZR1* in the rice bundle sheath could maintain chloroplast development without inducing premature senescence and decreased yield.

**Figure 4:**
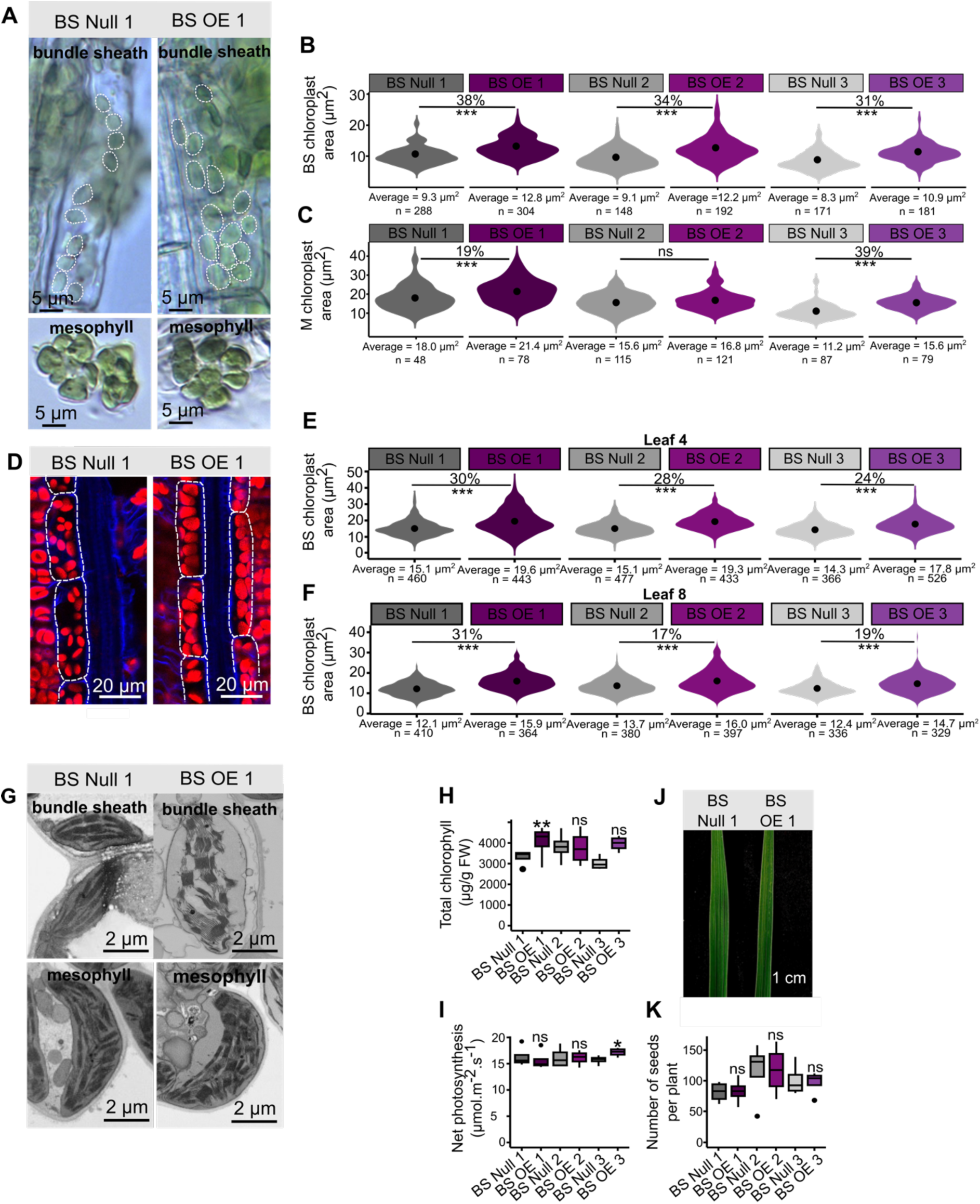
Bundle sheath cell-specific overexpression of *OsBZR1* increases the area of chloroplasts in rice bundle sheath cells with no impact on plant health. The rice codon optimised sequence for *BZR1* (rco*OsBZR1*) was cloned upstream of the rice bundle sheath cell-specific *SULPHITE REDUCTASE* promoter (p*OsSiR*) and transformed into rice to generate cell-specific overexpression lines (BS OE). **A:** Representative images from brightfield microscopy of individual bundle sheath and mesophyll cells from leaf 8. White dotted lines highlight individual chloroplasts within each bundle sheath cell. **B and C:** Chloroplast area in bundle sheath (BS) (**B**) and mesophyll (M) (**C**) cells from leaf 8. Chloroplast areas are from quantification of the brightfield microscopy. Percentages above violins indicate the change in chloroplast area of overexpressor lines compared with corresponding nulls. No statistically significant change in chloroplast area is represented by “ns”. The value below each violin is the mean chloroplast area calculated for that line and n represents the number of chloroplasts quantified. Four biological replicates were quantified for each line. **D:** Images derived from confocal laser scanning microscopy of bundle sheath cells and chloroplasts in leaf 4. Red and blue channels indicate chloroplasts and cell walls respectively. White dotted lines highlight bundle sheath cells. **E and F:** Bundle sheath cell chloroplast area in leaf 4 (**E**) and leaf 8 (**F**). Chloroplasts quantified are from the confocal microscopy. Percentages above violins indicate the change in chloroplast area of overexpressor lines compared with corresponding nulls. No statistically significant change in chloroplast area is represented by “ns”. The value below each violin is the mean chloroplast area calculated for that line and n represents the number of chloroplasts quantified. Four biological replicates were used for each line. **G:** Scanning electron microscope (SEM) images of bundle sheath and mesophyll cell chloroplasts from leaf 8. **H:** Total chlorophyll in leaf 8 of overexpressor lines compared with corresponding nulls. Data are from 4 biological repeats for each line. **I:** Rate of net photosynthesis under conditions of growth for overexpressor lines compared with corresponding nulls. Data are from 4 biological replicates for each line. **J:** Representative images of leaf 8 from null and overexpressor plants of the same age. **K:** Number of seeds produced by each line. Data are from 4 biological replicates for each line. For figures B, C, E, F, H, I, J and K stars above violins or boxes indicate a statistically significant difference between overexpressor and corresponding null as determined by independent t-test, where *p* ≤ 0.05 is flagged with one star (*), *p* ≤ 0.01 is flagged with 2 stars (**) and *p* ≤ 0.001 is flagged with three stars (***). No statistically significant change is represented by “ns”.

**Figure 5:**
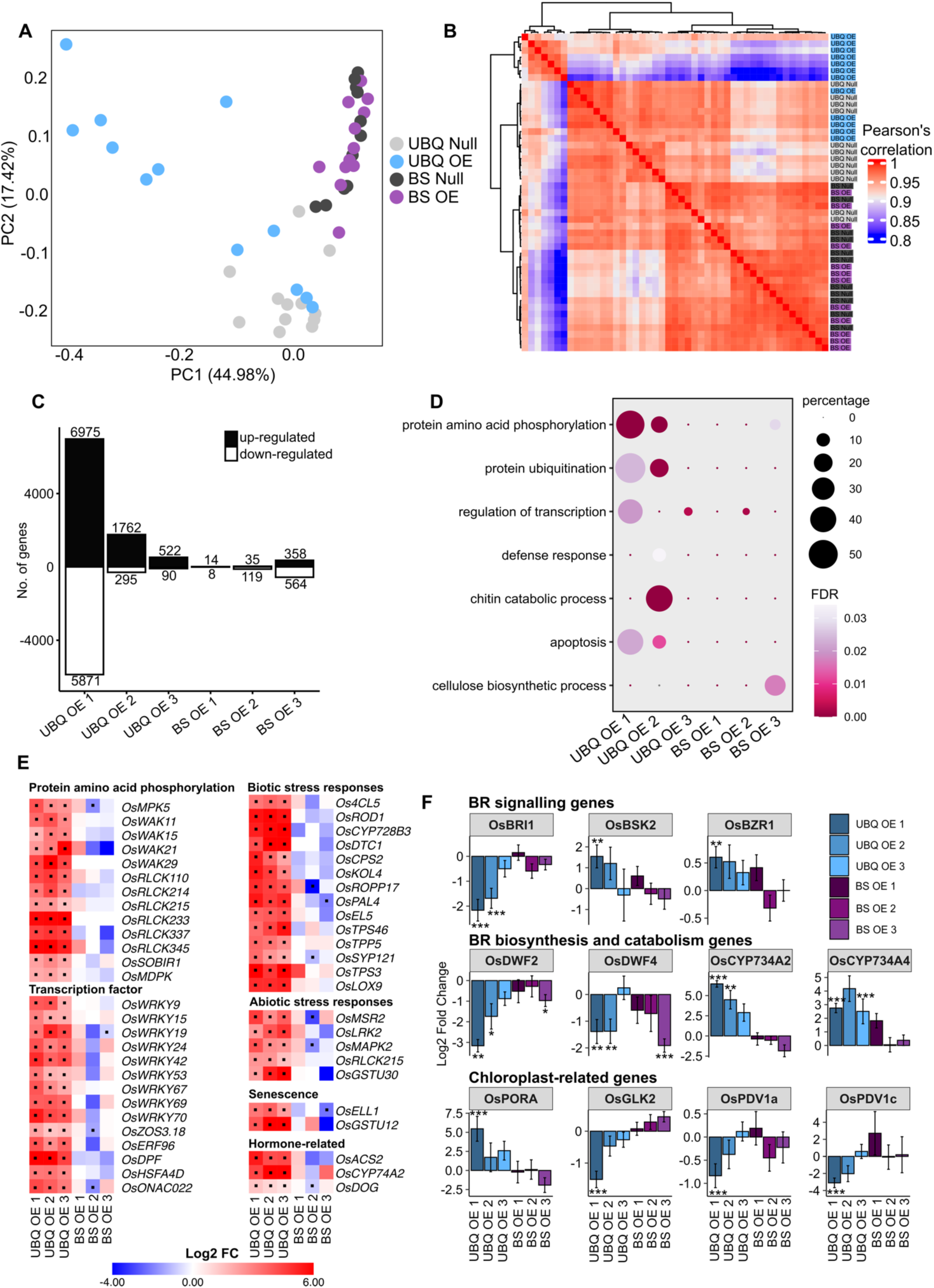
Transcriptome analyses of constitutive and bundle sheath cell-specific *OsBZR1* overexpressors. RNA from leaf 4 of 4 biological replicates from all of the UBQ Null, UBQ OE, BS Null and BS OE lines was used for cDNA library construction and subsequent transcriptome sequencing. **A:** Principal component analysis (PCA) indicates the transcriptome data separated primarily based on genotype. Different genotypes are denoted using different coloured circles. **B:** Heatmap showing relatedness of all samples based on Pearson’s correlation performed on log transformed data. **C:** Numbers of significantly (*p*-adj. < 0.05) differently expressed genes (DEGs) were determined by DESeq2 analysis which compared samples from an overexpressor with its corresponding null. Up-regulated genes are represented by black bars, and down-regulated genes by white bars. Numbers above each bar indicate the count of DEGs represented by the bar. **D:** Gene ontology (GO) terms enriched in the significantly differentially expressed gene lists from the overexpressors. Circle size depicts percentage, which is the number of genes in the given DEG list with the GO term divided by the number of genes in the reference genome with the GO term. The colour of each circle indicates the statistical significance of enrichment for each GO term, with FDR representing false discovery rate. **E:** Heatmaps showing the expression of genes of interest significantly differentially expressed in all three lines of constitutive overexpressors. The colour of each block indicates the Log2 fold change (FC) determined by DESeq2 analyses. A black dot within each block indicates a statistically significant change in Log2 FC. **F:** Log2 FC in expression of genes involved in brassinosteroid (BR) signalling, biosynthesis and catabolism and chloroplast development. Stars above bars indicate a statistically significant difference in Log2 FC expression where *p-*adj. ≤ 0.05 is flagged with one star (*), *p-*adj. ≤ 0.01 is flagged with 2 stars (**) and *p-*adj. ≤ 0.001 is flagged with three stars (***). Error bars represent standard error. Refer to Supp. dataset 5 for full gene names and gene IDs.

### Overexpression of OsBZR1 in the bundle sheath increases chloroplast area without inducing premature senescence

The promoter of the rice *SULPHITE REDUCTASE* (*SIR*) gene generates strong expression in bundle sheath cells (Hua et al., 2024) and so we used it to drive expression of *OsBZR1* (**Supp. fig. 8A**). Three independent homozygous and single copy overexpression lines (**Supp. fig. 8B**) along with corresponding null segregants were identified. RT-qPCR on T_2_ plants confirmed that the transgene was expressed in each line (**Supp. fig. 8C**). Hereafter these are referred to as BS Null 1, BS OE 1, BS Null 2, BS OE 2, BS Null 3 and BS OE 3. As with the constitutive overexpressor, individual mesophyll and bundle sheath cells from the BS Null and OE lines were isolated and brightfield microscopy used to quantify chloroplast planar area (**Fig. 4A, B and C**). All three overexpression lines contained larger bundle sheath cell chloroplasts when compared with corresponding null lines (**Fig. 4B**). Surprisingly, chloroplast areas in mesophyll cells were also increased compared with the corresponding nulls in two out of the three lines (**Fig. 4C**). Analysis of bundle sheath cells from leaf 4 and 8 by confocal laser scanning microscopy also indicated that chloroplasts in both cell types were larger in BS OE plants compared with nulls (**Fig. 4D, E & F**). No clear differences in ultrastructure between BS null and BS OE lines were discernible from scanning electron microscopy (**Fig. 4G**), nor could we detect differences in bundle sheath cell size (**Supp. fig. 9**).

Despite the statistically significant increase in bundle sheath chloroplast area, neither chlorophyll content nor rate of photosynthesis were consistently increased by cell-specific overexpression of *OsBZR1* (**Fig. 4H & I**). However, unlike the constitutive overexpressors, the leaves of the BS OE plants did not show premature senescence (**Fig. 4J**), and the number of seeds produced per plant did not differ from controls (**Fig. 4K**). Overall, these data confirm that increasing *OsBZR1* expression can increase chloroplast area in rice bundle sheath cells, but also that cell-specific perturbation avoided deleterious effects on growth.

### Constitutive overexpression of OsBZR1 perturbs stress-, brassinosteroid- and hormone-related pathways whilst bundle sheath cell-specific overexpression does not

To gain insight into how constitutive and cell-specific overexpression of *OsBZR1* re- programmed gene expression we conducted RNA sequencing on mature leaf 4 from UBQ Null, UBQ OE, BS Null and BS OE plants. Principal Component Analysis (PCA) showed that constitutive overexpression of *OsBZR1* caused the most variance and impacted on both the first and second components (**Fig. 5A**). In contrast, when *OsBZR1* was expressed in the bundle sheath there were few overall transcriptional changes, with samples from BS Null and BS OE clustering together (**Fig. 5A**). The trends in changes to transcript abundance were also evident from a heatmap derived from Pearson’s correlation analysis (**Fig. 5B**). Notably, correlation in mRNA abundance was weakest between UBQ OE and corresponding null lines whereas BS OE lines had high correlation with the corresponding BS null lines (**Fig. 5B**). Transcripts whose abundance was significantly different were identified by comparing each overexpression line with the corresponding null. Only genes with statistically significant changes (i.e. adjusted *p*-value < 0.05) were retained for subsequent analyses. This identified 6975, 1762 and 522 significantly up-regulated genes, and 5871, 295 and 90 genes down-regulated genes in the three constitutive overexpressors (**Fig. 5C, Supp. dataset 1 & 2**). Notably, the extent to which transcript abundance was perturbed corresponded to the degree of *OsBZR1* overexpression in these lines (**Fig. 5C, Supp. fig. 5C**). As would be expected from the PCA and Pearson’s correlation analysis, overexpression of *OsBZR1* from the bundle sheath promoter generated limited alterations to transcript abundance with only 14, 35 and 358 significantly up-regulated genes, and 8, 119 and 564 significantly down-regulated genes compared with null lines (**Fig. 5C, Supp. dataset 1 & 2**). As with the constitutive overexpressor, the extent to which transcript abundance was perturbed corresponded to the degree of *OsBZR1* overexpression in these lines (Fig. 5C, Supp. fig. 8C).

Gene Ontology (GO) enrichment analyses on transcripts that were up- and down- regulated identified several terms of interest, with the majority of these terms enriched in the constitutive overexpressing lines and not in the bundle sheath overexpressing lines (**Fig. 5D, Supp. dataset 3**). For example, GO terms for “protein amino acid phosphorylation”, “protein ubiquitination”, and “regulation of transcription” indicate a global change in transcription and protein degradation after constitutive *OsBZR1* overexpression. We note that several genes contributing to the GO terms involved in protein phosphorylation or transcription including four *Wall Associated Kinase* (*WAK*), six *Receptor-like Cytoplasmic Kinase* (*RLCK*) and nine *WRKY* transcription factors were significantly up-regulated in all three constitutive overexpressors, but this was not the case when *OsBZR1* expression was driven in the bundle sheath (**Fig. 5E, Supp. dataset 4**). Moreover, only the constitutive overexpressing line had enriched GO terms “defense response”, “chitin catabolic process” and “apoptosis” that are indicative of biotic and/or abiotic stress responses and are consistent with the increased senescence observed in the constitutive overexpressors. Specific genes associated with biotic and abiotic stress responses and senescence were also significantly up-regulated in constitutive overexpressors (**Fig. 5E**). Consistent with the lack of GO term enrichment in bundle sheath specific *OsBZR1* lines, transcript abundance of the genes involved in biotic and abiotic stress responses and senescence was not altered (**Fig. 5E, Supp. dataset 4**). It was notable that transcripts derived from the *OsACS2, OsCYP74A2* and *OsDOG* genes involved in ethylene, jasmonic acid and gibberellic acid synthesis were more abundant in all three constitutive *OsBZR1* overexpression lines, indicating possible hormonal crosstalk had been induced (**Fig. 5E, Supp. dataset 4**). Again, these genes were not differentially expressed in the lines when *OsBZR1* was driven from the bundle sheath promoter. Interestingly, the only enriched GO term specific to the bundle sheath overexpressor was “cellulose biosynthetic process” (**Fig. 5D**).

We next examined the expression of genes involved in brassinosteroid signalling, biosynthesis and catabolism to better understand how overexpression of *OsBZR1* perturbed these processes (**Fig. 5F, Supp. dataset 4**). In constitutive overexpressors, the brassinosteroid receptor *OsBRI1* was significantly down-regulated whilst the BR-signalling kinase *OsBSK2* and the endogenous *OsBZR1* gene were significantly up-regulated (**Fig. 5F, Supp. dataset 4**). This was not evident when *OsBZR1* was overexpressed in the bundle sheath. Moreover, two genes associated with brassinosteroid catabolism *OsCYP734A2* and *OsCYP734A4* were significantly up-regulated but only in the constitutive overexpressors (**Fig. 5F, Supp. dataset 4**). Additionally, transcripts from two brassinosteroid biosynthesis genes *OsDWF4* and *OsDWF2* were less abundant when *OsBZR1* was expressed constitutively and when expressed in the bundle sheath (**Fig. 5F, Supp. dataset 4**).

Finally, we interrogated the data to better understand how *OsBZR1* overexpression impacted genes involved in chloroplast function and biogenesis. We detected relatively few changes here, possibly due to the fact that the transcriptome of mature leaves was analysed at a stage during which chloroplast development may be complete. Transcripts derived from the chlorophyll biosynthesis gene *OsPORA* were more abundant in constitutive overexpressors, but this was not the case when *OsBZR1* was expressed in the bundle sheath (**Fig. 5F, Supp. dataset 4**). The *OsGLK2* transcription factor was significantly down- regulated in one of the constitutive overexpressing lines but slightly up-regulated when *OsBZR1* was overexpressed in the bundle sheath (**Fig. 5F, Supp. dataset 4**). Two genes involved in chloroplast division *PDV1a* and *PDV1c* were down-regulated when *OsBZR1* was constitutively overexpressed which could indicate the mechanism behind BZR1-modulated changes in chloroplast area (**Fig. 5F, Supp. dataset 4**).

## Discussion

### A role for brassinosteroids in controlling chloroplast size and number

As key phytohormones, brassinosteroids have been studied for decades with much of the work performed in *Arabidopsis thaliana*. One particularly well-documented role is the control of photomorphogenesis, including the induction of hypocotyl elongation combined with the inhibition of cotyledon expansion, chlorophyll biosynthesis and chloroplast differentiation in the dark (Asami et al., 2000; Chory et al., 1991; Komatsu et al., 2010; Tachibana et al., 2022; Yu et al., 2011; Zhang et al., 2021). Although it is known that brassinosteroids are important for chlorophyll accumulation and chloroplast gene expression during the transition from dark to light in multiple species, including rice, to our knowledge there are no reports demonstrating that brassinosteroids control chloroplast size or number. The results reported here show that exogenous treatment with an active brassinosteroid and a brassinosteroid biosynthesis inhibitor altered chloroplast planar area and number in rice bundle sheath cells and both constitutive and cell-specific overexpression of *OsBZR1* resulted in increased chloroplast area. Additionally, the transcript abundance of *OsPDV1a* and *OsPDV1c*, genes involved in chloroplast division, was significantly increased upon overexpression of *OsBZR1.* Together these results support a previously unknown role for brassinosteroids in modulating chloroplast size and number, potentially through the regulation of chloroplast division.

### Using OsBZR1 to engineer chloroplast volume specifically in rice bundle sheath cells

The role of BRs in increasing chloroplast area provided a new candidate to manipulate the chloroplast compartment of rice bundle sheath cells, an important characteristic if C_4_ photosynthesis is to be engineered into this species (Hibberd et al., 2008). Although constitutive overexpression of *OsBZR1* resulted in a significant increase in the area of individual chloroplasts in the bundle sheath, it did not impact mesophyll chloroplast area. This is possibly because the chloroplast compartment of mesophyll cells is already large, and so there is limited capacity to increase this further. Although the increased bundle sheath cell chloroplast area in the UBQ OE plants was extremely promising in terms of engineering this cell type, it had severe effects on plant health and yield. This is consistent with previous reports of misexpression of components of brassinosteroid biosynthesis or signalling leading to defects in plant growth (Clouse et al., 1996; Kim et al., 2020; Manghwar et al., 2022; Nolan et al., 2020). Exogenous application of brassinosteroids has been shown to have growth-promoting effects when a low dose is applied, whilst higher doses show growth retardation (Chaiwanon & Wang, 2015; González-García et al., 2011; Nolan et al., 2020). To our knowledge, the growth defects reported here are the first for rice and demonstrate conservation in the brassinosteroid signalling pathway between rice and Arabidopsis. Transcriptome analyses of the *OsBZR1* constitutive overexpressors showed differentially expressed genes with enriched GO terms including “biotic stress response”, “abiotic stress response” and “senescence”, consistent with the negative effects on plant development.

Endogenous tissue-specific control of brassinosteroid signalling ensures proper growth and avoids deleterious effects of constitutive brassinosteroid action (Nolan et al., 2020). Indeed, studies using tissue-specific promoters to complement brassinosteroid mutant phenotypes have revealed that cell type-specific confinement of brassinosteroid signalling is essential for proper shoot and root development (Chaiwanon & Wang, 2015; Hacham et al., 2011; Kang et al., 2017; Nolan et al., 2020, 2023; Savaldi-Goldstein et al., 2007; Vragović et al., 2015). This cell type-specific signalling can be harnessed for the development of stress resistant plants. For example, overexpression of BRL3, a vascular-enriched brassinosteroid receptor in Arabidopsis conferred drought stress tolerance without a growth penalty whereas altering the ubiquitously expressed BRI1 receptor conferred drought tolerance but at the expense of growth (Fàbregas et al., 2018). Although brassinosteroid signalling and regulatory mechanisms have been well elucidated in rice (Ahmar & Gruszka, 2022) this has not yet been defined at the level of single cells as is the case for Arabidopsis (Nolan et al., 2023), and so the results reported here confirm that endogenous tissue- specific control of brassinosteroid signalling is also present in this monocot crops.

As cell specific gene expression in Arabidopsis had proved useful (Fàbregas et al., 2018), we used a bundle sheath cell-specific promoter for rice (Hua et al., 2024) to express *OsBZR1* only in bundle sheath cells. This resulted in an increase in bundle sheath cell chloroplast area of up to 34% and had no detectable negative impacts on plant growth. Moreover, the transcriptome of these plants showed no enrichment in stress- or senescence-related GO terms, indicating that the cell-specific overexpression was successful in avoiding off-target perturbations. In fact, the transcriptome of these plants showed very little change compared with the corresponding nulls. A contributing factor to this outcome is likely that bundle sheath only makes up approximately 15% of chloroplast-containing leaf cells (Leegood, 2008). It was also noticeable that these lines showed little to no change in net photosynthesis or seed yield, again possibly due to the small proportion of bundle sheath cells in the leaf.

### Exploiting multiple master regulators to manipulate chloroplast development

The work presented here investigated the potential of manipulating brassinosteroid signalling to increase the chloroplast compartment in rice bundle sheath cells to mimic a more C_4_-like leaf anatomy. The percentage increases in bundle sheath chloroplast area of the constitutive and cell-specific *OsBZR1* overexpression lines reported here are comparable with other work describing master regulators of chloroplast development in rice. For example, overexpression of *ZmGLK* resulted in increases in chloroplast area of approximately 30% (Wang et al., 2017) and bundle sheath cell-specific overexpression of *OsCGA1* resulted in a 3.5-fold increase in chloroplast size and a significant increase in proportion of the bundle sheath cell occupied by chloroplasts (Lee et al., 2021). Based on these similarities in changes in chloroplast size, BZR1 could be considered as a promising new candidate for engineering the chloroplast compartment in rice. In the controlled conditions we used we observed no changes to photosynthesis or yield when *OsBZR1* was overexpressed, which contrasts with overexpression of *ZmGLK* that resulted in a 30-40% increase in vegetative biomass and grain yield in the field (Li et al., 2020). It may therefore be interesting to assess these *OsBZR1* overexpression lines in the field. In contrast to Arabidopsis, where BZR1 has been shown to modulate *GLK* expression (Yu et al., 2011) our transcriptome data showed no consistent changes to *OsGLK* when *OsBZR1* was overexpressed. Additionally, the expression of two other master regulators of chloroplast development, *OsCGA1* or *OsGNC*, was unchanged when *OsBZR1* was overexpressed. Thus, one could consider combining these regulators to test for additive or synergistic effects. To conclude, although work is still needed to elucidate precisely how brassinosteroids and BZR1 manipulate chloroplast size and number, the results shown here present a promising new candidate to be harnessed to manipulate chloroplast development in rice and, potentially, other important crop species.

## Materials and methods

### Plant material and growth conditions

For seed propagation and phenotyping experiments, seeds of wild-type (*Oryza sativa spp japonica* cv. Kitaake) and transgenic rice lines (UBQ Null, UBQ OE, BS Null and BS OE) were imbibed in sterile Milli-Q water and incubated at 28°C in the dark for two days. Seeds were transferred to Petri plates with moistened Whatman filter paper and germinated in the growth cabinet at 28°C with a 16/8-hour light/dark cycle for a further two days. Germinated seedlings were placed into 9 by 9 cm pots (two plants per pot) filled with Profile Field and Fairway soil amendment (www.rigbytaylor.com) and grown in a walk-in plant growth chamber under a 12-hour photoperiod at a photon flux density of 400 μmol m^-2^ s^-1^ at 28°C day and 20°C night. Plants were fed once a week with Peters Excel Cal-Mag Grower fertiliser solution (LBS Horticulture, Clone, UK) at a concentration of 0.33 g/L with additional iron (Fe7 EDDHA regular, Gardening Direct, UK) at a concentration of 0.065 g/L. Once fully expanded, leaf 4 and/or leaf 8 was harvested for phenotyping.

### Rice transformation

All constructs were generated using the Golden Gate cloning system (Engler et al., 2009; Weber et al., 2011). The full-length cDNA sequence of *OsBZR1* (LOC_Os07g39220) was rice codon optimised for ease of detection against the endogenous *OsBZR1* gene and domesticated to remove internal *Bpi*I and *Bsa*I restriction sites (**Supp. dataset 5**). For constitutive *OsBZR1* overexpression, a rice codon optimised *OsBZR1* sequence was cloned downstream of the maize *UBIQUITIN* promoter (p*ZmUBI*) and upstream of a nos terminator (nost) (**Supp. fig. 5**). The p*ZmUBI* was made up of 983 bp upstream of the transcription start site and 1014 bp of the first intron. For the bundle sheath cell-specific *OsBZR1* line, the rice codon optimised *OsBZR1* sequence was cloned downstream of the rice *SULFITE REDUCTASE* promoter (p*OsSIR*) (Hua et al., 2024) and upstream of a nos terminator (nost) (**Supp. fig. 8**). These level 1 modules were then cloned with a hygromycin resistance level 1 module which contained the hygromycin resistance gene downstream of the rice *ACTIN* promoter (p*OsACT)* and upstream of a nos terminator. This module was used for selection of transformants at T_0_ stage and selection of homozygous lines at T_2_ stage.

*Oryza sativa spp. japonica* cv. Kitaake was transformed using *Agrobacterium tumefaciens* as described previously (Hiei & Komari, 2008) with several modifications. Seeds were de-husked and sterilized with 10% (v/v) bleach for 15 min before placing them on nutrient broth (NB) callus induction media containing 2 mg/L 2,4-dichlorophenoxyacetic acid for 4 weeks in the dark at 28°C. Calli were co-incubated with *A. tumefaciens* strain LBA4404 carrying the expression plasmid of interest in NB inoculation medium containing 40 μg/ml acetosyringone for 3 days in the dark at 22°C. Calli were transferred to NB recovery medium containing 300 mg/L timentin for 1 week in the dark at 28°C. They were then transferred to NB selection medium containing 35 mg/L hygromycin B for 4 weeks in the dark at 28°C. Proliferating calli were subsequently transferred to NB regeneration medium containing 100 mg/L myo-inositol, 2 mg/L kinetin, 0.2 mg/L 1-naphthaleneacetic acid, and

0.8 mg/L 6-benzylaminopurine for 4 weeks in the light at 28°C. Plantlets were transferred to NB rooting medium containing 0.1 mg/L 1-naphthaleneacetic acid and incubated in Magenta pots for 2 weeks in the light at 28°C. Finally, plants were transferred Profile Field and Fairway soil amendment (www.rigbytaylor.com) and grown in a walk-in plant growth chamber under a 12-hour photoperiod at a photon flux density of 400 μmol m^-2^ s^-1^ at 28°C day and 20°C night. DNA was isolated from individual T_0_ plants and DNA blot analysis performed to determine insertion copy number. Lines with single insertions in different locations of the genome were used for phenotyping experiments.

### BL and Brz treatments and chlorophyll analysis

*Oryza sativa spp. japonica* cv. Kitaake seeds were de-husked and sterilized in 10% (v/v) bleach for 30 min. After washing several times with sterile water, seeds were imbibed in water and incubated at 28°C in the dark for two days. For de-etiolation experiments, germinated seedlings were transferred in a dark room equipped with a green light into ½ strength Murashige and Skoog (MS) medium (0.8% agar) supplemented with either 10 µM brassinolide (Santa Cruz Biotechnology, Inc.) or 10 µM brassinazole (Merck Life Science UK Ltd., Gillingham, UK). Magentas were covered in aluminium foil and placed in a growth cabinet set to a 28°C, 16-hour day and 20°C, 8-hour night cycle for a further three days. At the beginning of the fourth photoperiod, aluminium foil was removed from the magentas to expose the seedlings to light. Leaf tissue was harvested for chlorophyll quantification and confocal microscopy at 0, 4, 12 and 24 hours after exposure to light. For later stages, sterile germinated seedlings were transferred into ½ strength Murashige and Skoog medium (0.8% agar) supplemented with either 10 µM Brassinolide or 10 µM Brassinazole and placed in a growth cabinet set to a 28°C, 16-hour day and 20°C, 8-hour night cycle until leaf 4 had fully expanded, at which time it was harvested for chlorophyll quantification and confocal microscopy.

Tissue for chlorophyll quantification (leaf 4, leaf 8 or seedlings during de-etiolation experiments) was harvested, weighed and immediately flash-frozen in liquid nitrogen. Frozen tissue was ground into a fine powder and suspended in 1 ml of 80% (v/v) acetone.

After vortexing, the tissue was incubated on ice for 15 min with occasional mixing of the suspension. The tissue was spun at 13 000 rpm for 5 min at 4°C and supernatant removed. The extraction was repeated, and supernatants pooled before measuring absorbance at 663.6 nm and 646.6 nm. Total chlorophyll content was determined as described previously (Porra et al., 1989).

### Chloroplast imaging and quantification

Chloroplasts in individual bundle sheath and mesophyll cells were imaged and quantified using light and confocal laser microscopy. To isolate single mesophyll and bundle sheath cells for light microscopy, cells from leaf 8 of wild-type Kitaake or transgenic rice lines (UBQ Null, UBQ OE, BS Null and BS OE) were isolated following the protocol of Khoshravesh & Sage (2018). Briefly, the middle region of fully expanded leaf 8 was cut into 5 mm-long x 2 mm-wide strips along the leaf proximodistal axis using a razorblade and immediately immersed in room temperature 4% w/v paraformaldehyde (pH 6.9) (Thermo Fisher Scientific Inc.). Fixed tissue was left in paraformaldehyde at 4°C for at least one hour (up to overnight). Cell walls were then digested by incubating in 0.2M sodium-EDTA (pH 9.0) at 55°C for 2 hours, rinsed in digestion buffer (0.15M Na_2_HPO_4,_ 0.04M citric acid, pH 5.3) and then incubated in 2% w/v pectinase from *Aspegillus niger* (Merck Life Science UK Ltd., Gillingham, UK) in digestion buffer at 45°C for 2 hours. Digestion was stopped by incubation in empty digestion buffer twice for 30 minutes at room temperature. After digestion, individual cells were released by mechanical disruption using the bottom of an Eppendorf tube. Isolated mesophyll and bundle sheath cells were imaged by brightfield microscopy using an Olympus BX41 microscope (Olympus UK and Ireland, Southend-on-Sea, UK), recording each cell in the paradermal plane where most of the chloroplasts were in focus. Images were captured using an MP3.3-RTV-R-CLR-10-C MicroPublisher camera and QCapture Pro 7 software (Teledyne Photometrics, Birmingham, UK). The area of individual mesophyll and bundle sheath cell chloroplasts in each image was quantified using ImageJ version 1.53k.

To visualise and quantify chloroplasts in bundle sheath cells using a confocal laser microscope, the methods described in Billakurthi and Hibberd (2023) were followed. Briefly, the middle region of fully expanded leaf 4 and 8 was fixed with 1% (w/v) glutaraldehyde (Thermo Fisher Scientific Inc.) in 1X PBS buffer. Samples were left in fixative for two hours and then washed twice with 1X PBS buffer. Before confocal microscopy, the adaxial side of the fixed leaf material was ablated gently with a fine razor blade to remove mesophyll layers and then incubated in calcofluor white (0.1%; Sigma) for 5 mins to stain cell walls prior to rinsing twice with H_2_O. A Leica SP8X confocal microscope upright system (Leica Microsystems) was used for fluorescence imaging of the bundle sheath cell chloroplasts and cell walls. Imaging was conducted using a 25X water immersion objective and Leica Application Suite X software (LAS X; version: 3.5.7.23225). Calcofluor white was excited at 405 nm and emitted fluorescence captured from 452 to 472 nm. Chlorophyll autofluorescence was excited at 488 nm and emission captured 672 to 692 nm. For all lines, leaf 4 and 8 from 4 plants were assessed, with 3 different intermediate veins imaged in each leaf. The planar area of individual mesophyll and bundle sheath chloroplasts in each image was quantified using ImageJ version 1.53k.

### Serial block-face scanning electron microscopy

To visualise the ultrastructure of individual chloroplasts, serial block-face scanning electron microscopy (SEM) was used. For this, the middle region of fully expanded leaf 8 was cut into 2 mm x 2 mm squares and fixed in 2% (v/v) glutaraldehyde and 2% (w/v) formaldehyde in 0.05 - 0.1 M sodium cacodylate (NaCac) buffer (pH 7.4) containing 2 mM calcium chloride. Samples were vacuum infiltrated overnight, washed 5 times in 0.05 – 0.1 M NaCac 557 buffer, and post-fixed in 1% (v/v) aqueous osmium tetroxide, 1.5% (w/v) potassium ferricyanide in 0.05 M NaCac buffer for 3 days at 4°C. After osmication, samples were washed 5 times in deionized water and post-fixed in 0.1% (w/v) thiocarbohydrazide for 20 min at room temperature in the dark. Samples were then washed 5 times in deionized water and osmicated for a second time for 1 hr. in 2% (v/v) aqueous osmium tetroxide at room temperature. Samples were washed 5 times in deionized water and subsequently stained in 2% (w/v) uranyl acetate in 0.05 M maleate buffer (pH 5.5) for 3 days at 4°C and washed 5 times afterwards in deionized water. Samples were then dehydrated in an ethanol series, transferred to acetone, and then to acetonitrile. Leaf samples were embedded in Quetol 651 resin mix (TAAB Laboratories Equipment Ltd.) and cured at 60°C for 2 days. Ultra-thin sections of embedded leaf samples were prepared and placed on Melinex (TAAB Laboratories Equipment Ltd) plastic coverslips mounted on aluminium SEM stubs using conductive carbon tabs (TAAB Laboratories Equipment Ltd), sputter-coated with a thin layer of carbon (∼30 nm) to avoid charging and imaged in a Verios 460 scanning electron microscope at 4 keV accelerating voltage and 0.2 nA probe current using the concentric backscatter detector in field free (low magnification) or immersion (high magnification) mode (working distance 3.5 – 4 mm, dwell time 3 μs, 1536 x 1024 pixel resolution). SEM stitched maps were acquired at 10,000X magnification using the FEI MAPS automated acquisition software. Greyscale contrast of the images was inverted to allow easier visualisation.

### Gas exchange measurements

Fully expanded leaf 8 was used to measure photosynthetic rates using a LI-6800 photosynthesis system (LICOR Biosciences). For the UBQ OE plants, measurements were performed prior to the onset of early senescence. Four individual plants were sampled per line. Measurements were made at a constant airflow of 400 μmol.s^-1^, leaf temperature of 30°C and relativity humidity of 60%. Leaves were acclimated in the chamber for approximately 10 mins before net photosynthesis (CO_2_ assimilation rate - A) measurements were made at ambient conditions (light intensity of 400 μmol photons m^−2^ s^−1^ and intercellular CO_2_ concentration (Ci) of 400 μmol.CO_2_ mol^−1^ air). All measurements were performed on the midportion of the lead blade.

### Phylogenetic tree inference

To identify rice orthologs of transcription factors that mediate BR-responsive gene expression in Arabidopsis, the protein coding genes derived from representative gene models were downloaded from Phytozome (Goodstein et al., 2012). These proteomes were subject to orthogroup inference using OrthoFinder (Emms & Kelly, 2015, 2019). The orthogroup containing Arabidopsis BZR1, BES1, (BZR2), BEH1, BEH2, BEH3 and BEH4 was identified. The sequences from this orthogroup were subject to multiple sequence alignment using MergeAlign (Collingridge & Kelly, 2012) followed by bootstrapped maximum likelihood phylogenetic tree inference using IQTREE2 (Minh et al., 2020) with the best fitting model of sequence evolution (JTT+I+G4) inferred from the data.

### Total RNA extraction, cDNA library preparation and transcriptome analysis

Total RNA was extracted from leaf 4 using the RNeasy Plant Mini Kit (Qiagen, Germany) according to the manufacturer’s instructions. Genomic DNA was removed from each sample using the RNase-Free DNase Set (Qiagen, Germany). RNA degradation and contamination was checked on 1% (w/v) agarose gels, then RNA quality and concentration determined with the RNA600 Pico Assay using the Agilent 2100 Bioanalyzer (Agilent Technologies, USA) and ND-200 nanodrop (NanoDrop Technologies Inc., USA). RNA was sent to the Novogene Genomic Sequencing Centre (Cambridge) for library preparation and sequencing using the Illumina NovaSeq 600 PE150 sequencing platform and strategy. At least 6 Gb of raw data per sample was generated for subsequent transcriptome analysis.

Quality of raw sequencing data was assessed and controlled using the FastQC platform version 0.11.4 (Andrew, S. 2010). Adapter trimming and filtering of all low quality reads was performed using BBDuk (https://www.geneious.com/plugins/bbduk/) with the following parameters: k=13, ktrim=r, useshortkmers=t, mink=5, qtrim=r, trimq=2- minlength=50, ftl=10, ftr=139. The *Oryza sativa* IRGSP-1.0 transcriptome was downloaded from Ensemble Plants (https://plants.ensemble.org) and used to build a Salmon reference index which was subsequently used to quantify the cleaned reads (Salmon version 1.5.2). For Salmon quantification (Patro et al., 2017), all parameters were left as default. The Salmon alignment and quantification results were checked using MultiQC (Ewels et al., 2016). To check that biological replicates clustered together and to visualise how the overexpression lines differed from corresponding null lines and from each other, a principal component analysis (PCA) was performed using the ggfortify package in R. Transcripts per million (TPM) counts from Salmon alignments were filtered for genes with at least 48 counts across all samples. The filtered data were then normalised to account for library size and transformed to a log scale using rlog transformation to allow for easier visualisation. For the Pearson’s correlation heatmap, the filtered, rlog transformed TPM data was used.

To determine changes in expression of individual genes between OE lines and corresponding null lines, the DESeq2 package (version 4.2) in R was used (Love, Huber & Anders, 2014). The quant.sf file generated for each sample from the Salmon quantification was used as the input for the DESeq2 analysis. Independent DESeq2 analyses were performed for the two different overexpression lines (i.e. UBQ OE samples versus corresponding UBQ Null samples and BSC OE samples versus corresponding BSC Null samples). Lists of significantly differentially expressed genes for further investigation were developed by filtering for genes with an adjusted *p*-value less than 0.05. To gain further biological insight into DEG lists, Gene Ontology (GO) enrichment analyses were performed using AgriGO v2 (Tian et al., 2017) with the *Oryza sativa* MSU7.0 genome set as the background reference.

### Quantitative real time PCR (RT-qPCR)

To check transgene expression levels, RNA was extracted from fully expanded leaf 4 as described above. The total RNA was used as a template to synthesize cDNA using the SuperScript II Reverse Transcriptase kit (Invitrogen) according to the manufacturer’s instructions. RT-qPCR was carried out on the cDNA using SYBR Green JumpStart Taq ReadyMix (Merck Life Science UK Ltd., Gillingham, UK) on a CFX384 Real-Time System (Bio-Rad) with the following cycle parameters; 94°C for 2 min, (94°C for 15 sec, 60°C for 1 min) x 40. This was performed using *OsBZR1* specific primers and *OsEF-1*⍺ and *OsUBQ* reference genes (Jain et al., 2006; Jain et al., 2018) with the following sequences.

*OsBZR1-*F: CGTACAACCTCGTGAACCC,

*OsBZR1-*R: CGTCACCCTACCTTTGTCG,

*OsEF-1⍺-*F: TTTCACTCTTGGTGTGAAGCAGAT,

*OsEF-1⍺-*R: GACTTCCTTCACGATTTCATCGTAA,

*OsUBQ5*-F: ACCACTTCGACCGCCACTACT,

*OsUBQ5*-R: ACGCCTAAGCCTGCTGGTT

### Data analyses

Unless otherwise stated, all statistical analyses were performed using StatSoft Statistica software.

## Funding and acknowledgements

The work was funded by BBSRC grant BBP0031171 to J.M.H. S.K. was supported by a Royal Society University Research Fellowship. Work in S.K.’s laboratory was supported by the European Union’s Horizon 2020 research and innovation programme under grant agreement no. 637765 and by the Wellcome Trust under grant number 226598/Z/22/Z.

R.W.H. was supported by a BBSRC studentship through BB/J014427/1. We acknowledge Dr. Karin Müller from the Cambridge Advanced Imaging Centre and Dr. Tina Shreier for their significant help with SEM imaging.

## Author contributions

S.K. and R.W.H. conducted the phylogenetic analysis and identified BZR1 as a potential regulator of chloroplast biogenesis. R.W.H designed the coding sequences for the overexpression and RNAi constructs. A.R.G.P cloned the DNA constructs for stable rice transformation. S.S carried out the stable rice transformation. L.H.L grew and genotyped the constitutive overexpression lines and contributed intellect and discussion throughout the project. L.C. conducted all experiments and analysed the data. L.C. and J.M.H wrote the manuscript with input from all authors. J.M.H guided execution of experiments and oversaw the project.

## Supplementary Figure legend titles

**Supplemental Figure 1:** Bundle sheath cell size of untreated, BL and Brz treated seedlings during de-etiolation.

**Supplemental Figure 2:** Number of veins in leaf 4 from untreated, BL and Brz treated plants.

**Supplemental Figure 3.** Maximum likelihood phylogenetic tree of the BZR1 gene family in plants.

**Supplemental Figure 4:** Development of *OsBZR1* knockout lines through RNA interference (RNAi).

**Supplemental Figure 5:** Development of rice codon optimised *OsBZR1* constitutive overexpressing lines (UBQ OE).

**Supplemental Figure 6:** Total chlorophyll content in leaf 8 of UBQ Null and UBQ OE lines.

**Supplemental Figure 7:** Bundle sheath cell size in UBQ Null and UBQ OE lines.

**Supplemental Figure 8:** Development of rco*OsBZR1* bundle sheath cell-specific overexpressing lines.

**Supplemental figure 9:** Bundle sheath cell size in BS Null and BS OE lines.

