## Supplemental figures for "Increased chloroplast area in the rice bundle sheath through cell specific perturbation of brassinosteroid signalling"

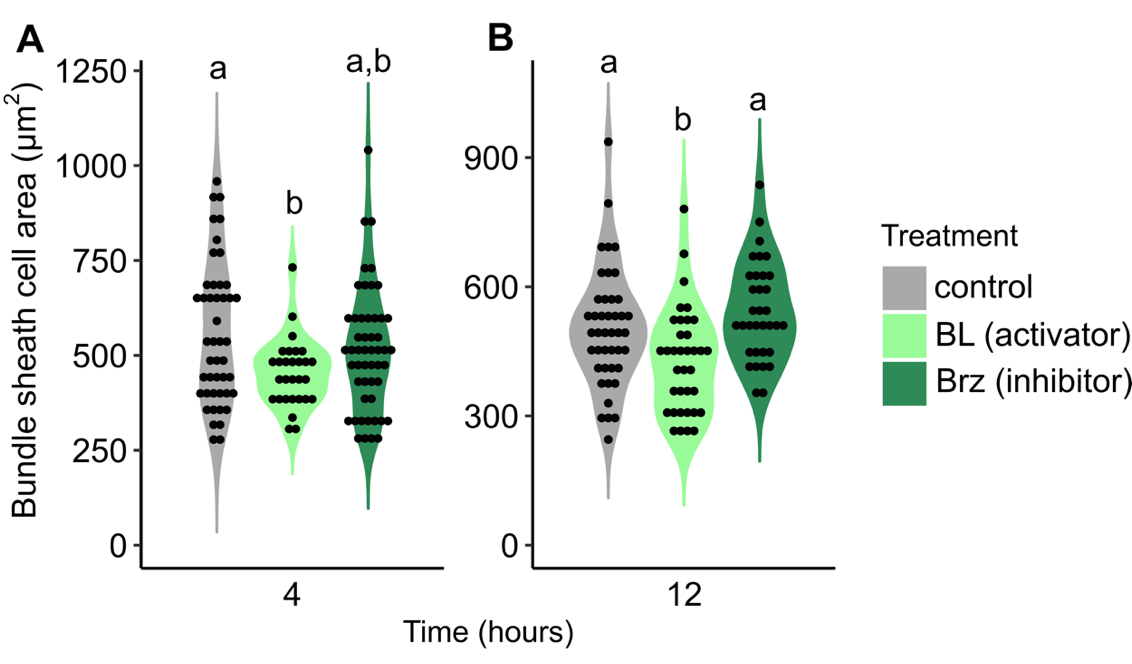

**Supplemental Figure 1: Bundle sheath cell size of untreated, BL and Brz treated seedlings during de-etiolation.** Seeds were germinated in water and transferred in the dark to ½ MS-agar media with or without 10 µM brassinolide (BL) or 10 µM brassinazole (Brz). After 4 days seedlings were transferred to light and shoot tissue harvested 0, 4, 12 and 24 hours later for chlorophyll quantification and imaging using confocal laser scanning microscopy. **A** and **B:** Bundle sheath cell area in control and BL/Brz treated seedlings 4 (A) and 12 (B) hours after exposure to light. Data are derived from confocal microscopy and from at least 30 cells for each timepoint in each treatment. Letters above violins represent statistically significant differences (*p* ≤ 0.05) in mean values as determined by Fisher LSD post-hoc analysis following a one-way ANOVA.

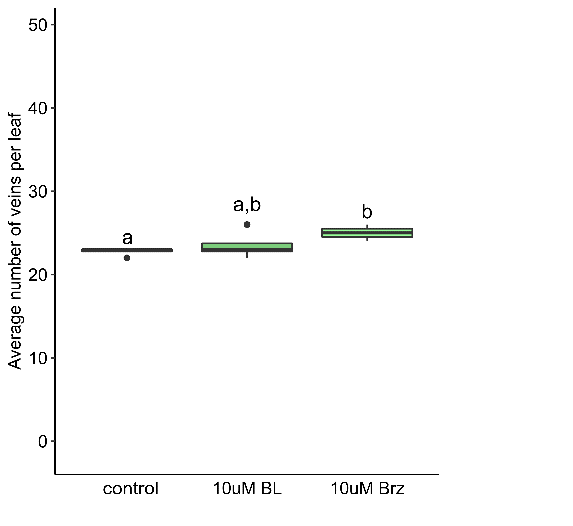

**Supplemental Figure 2: Number of veins in leaf 4 from untreated, BL and Brz treated plants.** Seeds were germinated in water and transferred to untreated MS media or MS media supplemented with 10 µM brassinolide (BL) or 10 µM brassinazole (Brz) and the number of veins in fully expanded leaf 4 counted. Data are from 4 leaves for each treatment. Letters above boxes represent statistically significant differences (*p* ≤ 0.05) in mean values as determined by Fisher LSD post-hoc analysis following a one-way ANOVA.

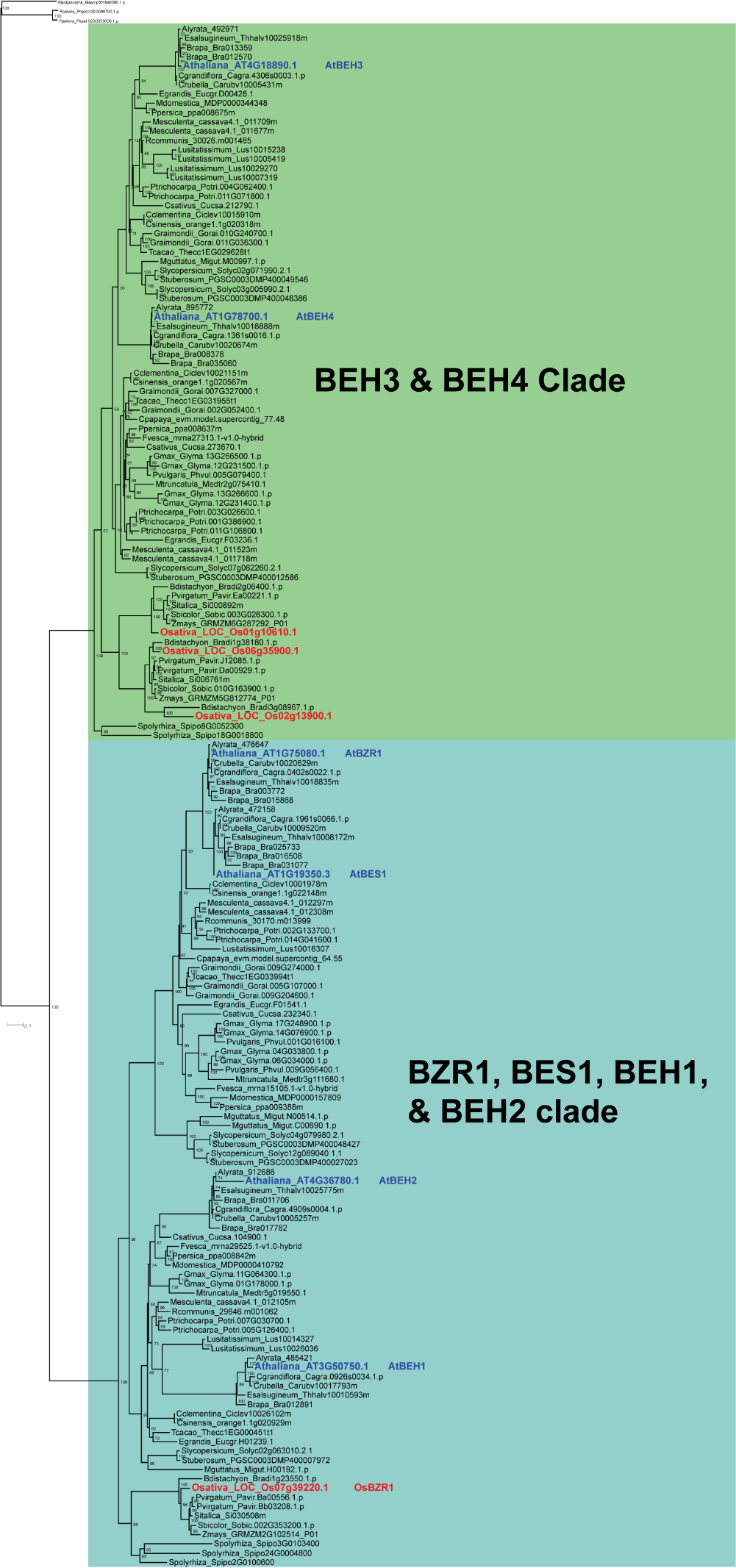

**Supplemental Figure 3. Maximum likelihood phylogenetic tree of the BZR1 gene family in plants.** *Arabidopsis thaliana* genes are highlighted in blue font with gene names added after the accession number. *Oryza sativa* accession numbers are highlighted in red font with *OsBZR1* indicated on the tree. Bootstrap support values shown at internal nodes. Scale bar indicates number of substitutions per aligned sequence site.

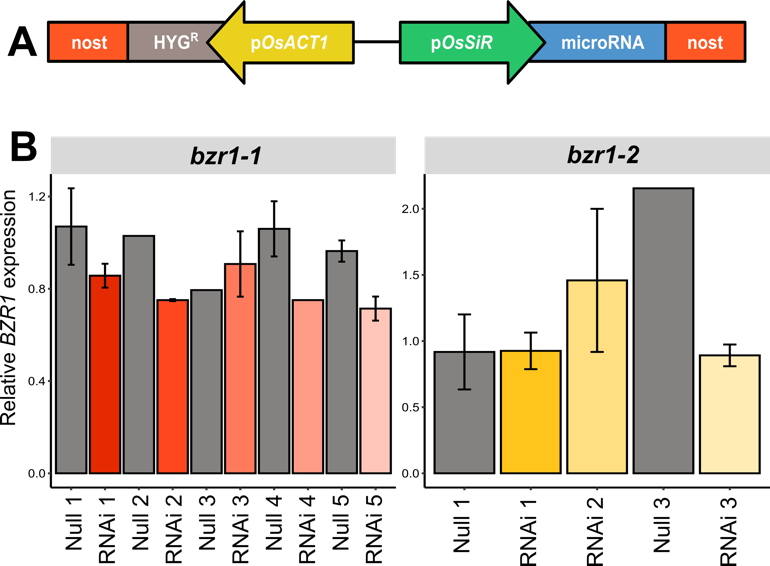

**Supplemental Figure 4: Development of *OsBZR1* knockout lines through RNA interference (RNAi). A:** Schematic of the construct used for transformation. The hygromycin resistance gene (HYG^R^) driven by the ACTIN promoter (p*OsACT1*) was used to select transformants. The maize UBIQUITIN promoter (p*ZmUBI1*) was used to drive *OsBZR1* specific microRNA sequences for repression of expression. **B:** Homozygous lines in the T_2_ generation were identified and the level of *OsBZR1* overexpression in leaf 4 determined by RT-qPCR. Expression is shown relative to *OsUBI* reference gene and represents the mean derived from one to four biological replicates. Error bars represent standard errors. The two different lines (*bzr1-1* and *bzr1-2*) were generated using different microRNA sequences.

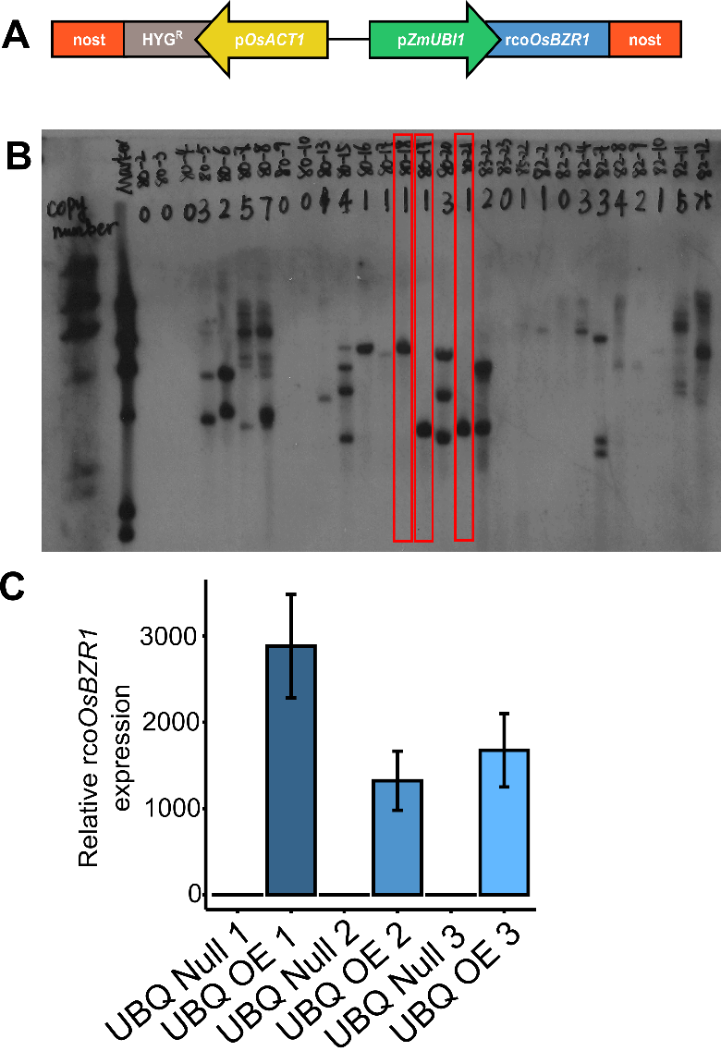

**Supplemental Figure 5: Development of rice codon optimised *OsBZR1* constitutive overexpressing lines (UBQ OE).** **A:** Schematic of the construct used for transformation. The hygromycin resistance gene (HYG^R^) driven by the ACTIN promoter (p*OsACT1*) was used to select transformants. The maize UBIQUITIN promoter (p*ZmUBI1*) was used to drive constitutive expression of rice codon optimised *BZR1* (rco*OsBZR1*). **B:** Southern blot performed on DNA from T_0_ transformants to identify lines containing single copies of the T-DNA insert. Red boxes outline the three independent, single copy lines chosen for phenotyping. 80-18, 80-19 and 80-21 refer to UBQ OE 1, UBQ OE 3 and UBQ OE 2 respectively **C:** Homozygous lines in the T_2_ generation were identified and the level of rco*OsBZR1* overexpression in leaf 4 determined by RT-qPCR. Expression is shown relative to *OsEF1α* and *OsUBI* reference genes and represents the mean derived from four biological replicates. Error bars represent standard errors.

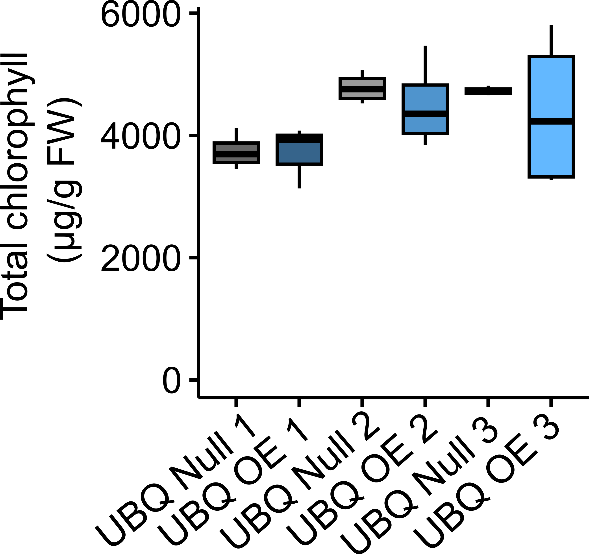

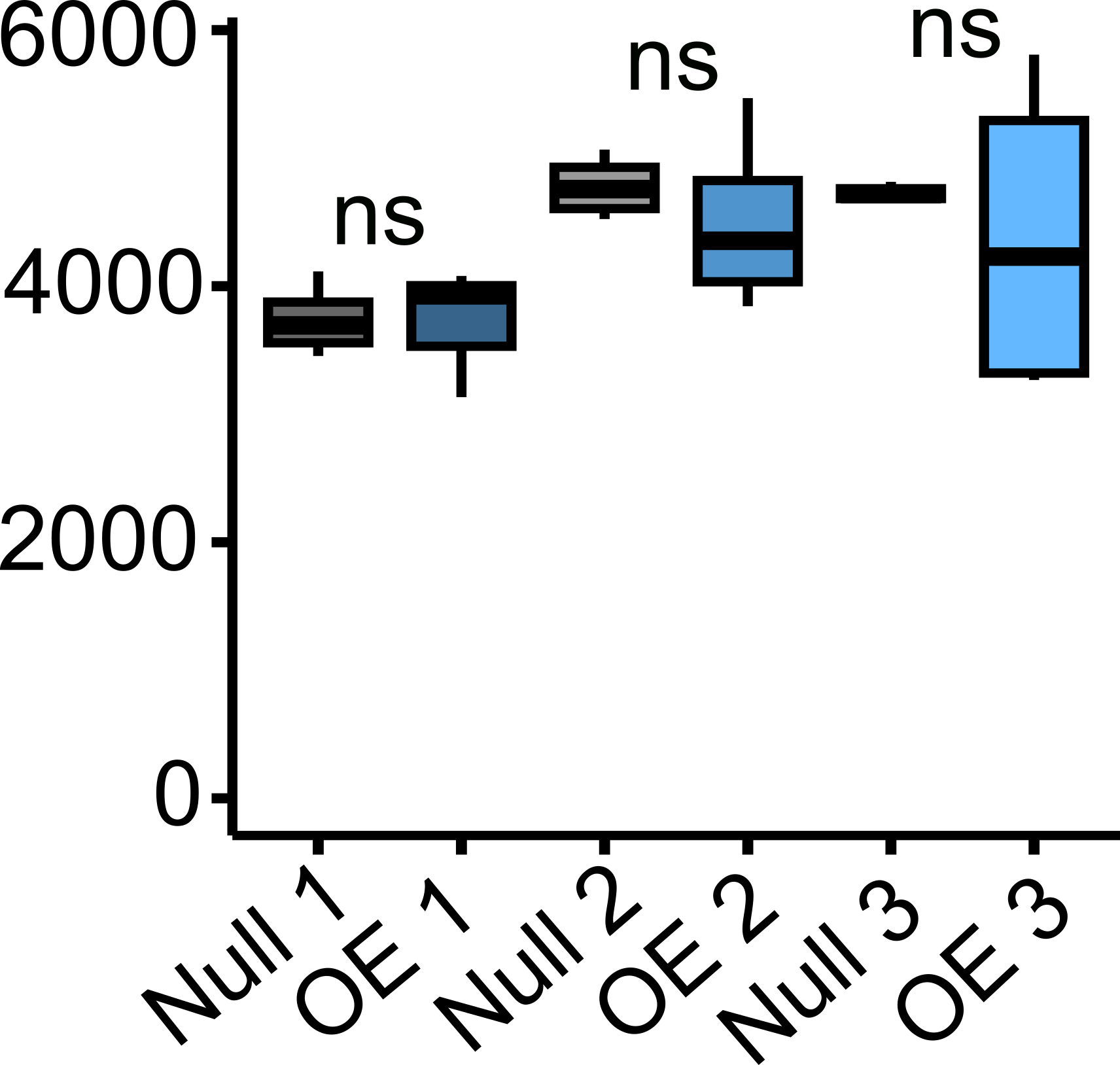

**Supplemental Figure 6: Total chlorophyll content in leaf 8 of UBQ Null and UBQ OE lines.** The rice codon optimised sequence for *BZR1* (rco*OsBZR1*) was cloned upstream of the maize *UBIQUITIN* promoter (p*ZmUBI*) and transformed into Kitaake rice to generate constitutive overexpression lines (referred to as UBQ OE). Total chlorophyll (µg/g fresh weight (FW)) from fully expanded leaf 8 was determined. Data are from 4 biological repeats for each treatment. No statistically significant change in chlorophyll content between overexpressor and corresponding null is represented by “ns” as determined by independent t-test.

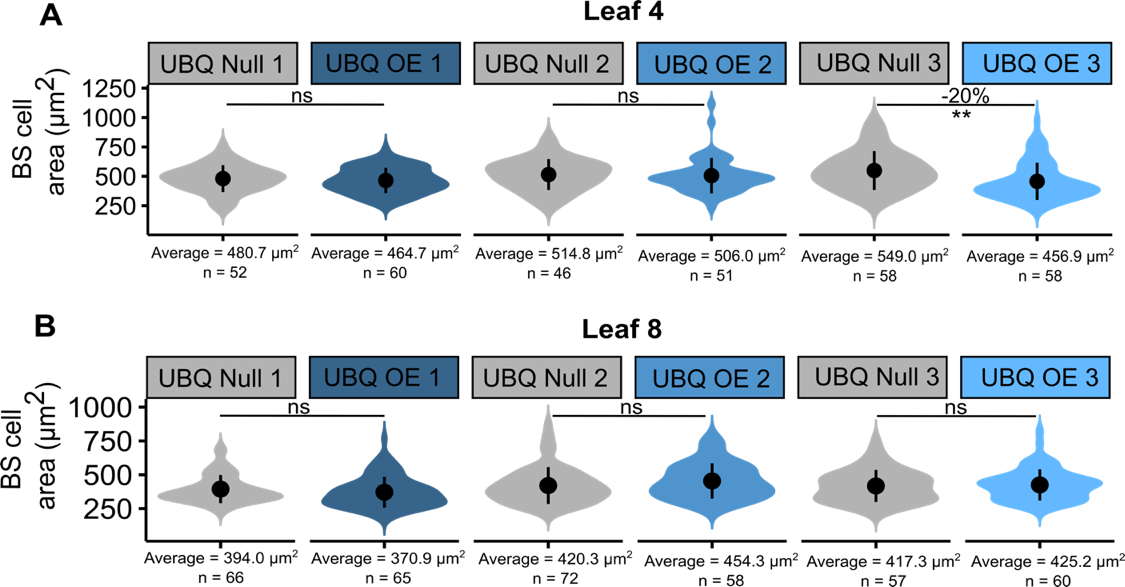

**Supplemental Figure 7:** **Bundle sheath cell size in UBQ Null and UBQ OE lines.** The rice codon optimised sequence for *BZR1* (rco*OsBZR1*) was cloned upstream of the maize *UBIQUITIN* promoter (p*ZmUBI*) and transformed into Kitaake rice to generate constitutive overexpression lines (referred to as UBQ OE). The area of individual bundle sheath (BS) cells in leaf 4 (**A**) and leaf 8 (**B**) of UBQ Null and UBQ OE plants was calculated from confocal microscope images. Percentage values above the violins indicate the change in BS cell area of UBQ OE chloroplasts compared to the corresponding null line. No statistically significant change in BS cell area is represented by “ns”. The average below each violin is the average BS cell area calculated for that line and n represents the number of BS cells quantified for this average. Four biological replicates were used for each line. Stars above violins or boxes indicate a statistically significant difference between UBQ OE and corresponding null lines as determined by independent t-test, where *p* ≤ 0.05 is flagged with one star (*), *p* ≤ 0.01 is flagged with 2 stars (**) and *p* ≤ 0.001 is flagged with three stars (***).

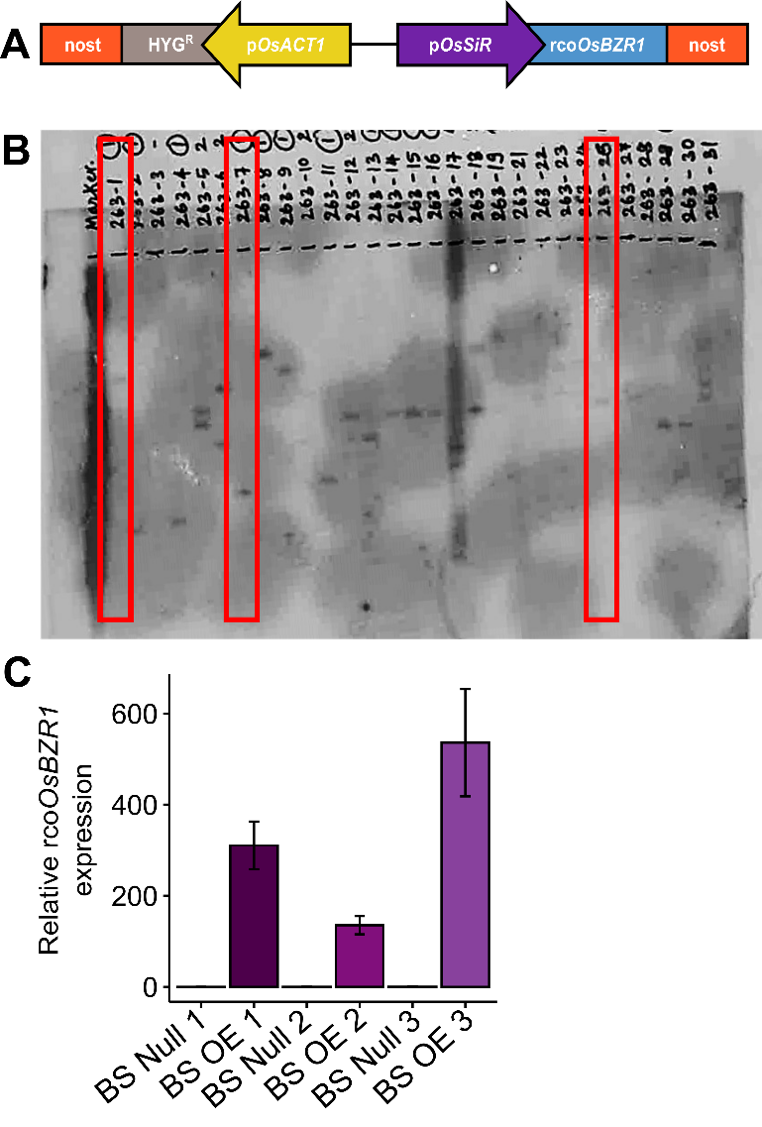

**Supplemental Figure 8:** **Development of** **rco*OsBZR1* bundle sheath cell-specific overexpressing lines.** **A:** Schematic of the construct used for transformation. The hygromycin resistance gene (HYG^R^) driven by the ACTIN promoter (p*OsACT1*) was used to select transformants. The rice SULPHITE REDUCTASE promoter (p*OsSiR*) was used to drive bundle sheath cell-specific expression of rice codon optimised BZR1 (rco*OsBZR1*). **B:** Southern blot performed on DNA from T_0_ transformants to identify lines containing single copies of the T-DNA insert. Red boxes outline the three independent, single copy lines chosen for phenotyping. 263-1, 263-26 and 263-7 refer to BS OE 1, BS OE 3 and BS OE 2 respectively. **C:** Homozygous lines in the T_2_ generation were identified and the level of rco*OsBZR1* overexpression in leaf 4 determined by RT-qPCR. Expression is shown relative to *OsEF1α* and *OsUBI* reference genes and the mean from four biological replicates presented. Error bars represent standard errors.

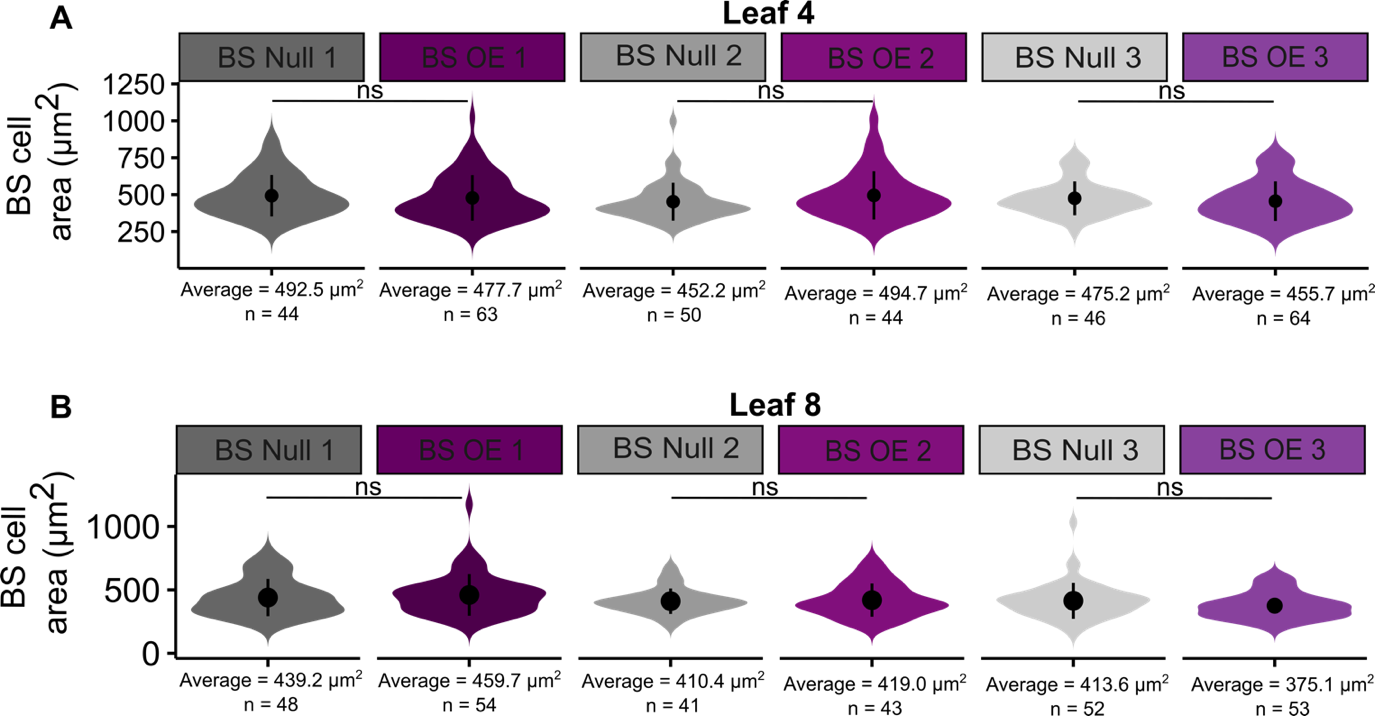

**Supplemental figure 9:** **Bundle sheath cell size in BS Null and BS OE lines.** The rice codon optimised sequence for *BZR1* (rco*OsBZR1*) was cloned upstream of the rice bundle sheath cell-specific *SULPHITE REDUCTASE* promoter (p*OsSIR*) and transformed into Kitaake rice to generate cell-specific overexpression lines (referred to as BS OE). The area of individual bundle sheath (BS) cells in leaf 4 (**A**) and leaf 8 (**B**) of BS Null and BS OE plants was determined from confocal microscopy. Percentage values above the violins indicate the change in BS cell area of BS OE chloroplasts compared with corresponding null lines. No statistically significant change in BS cell area is represented by “ns”. The average below each violin is the average BS cell area calculated for that line and n represents the number of BSs quantified. Four biological replicates were used for each line. Stars above violins or boxes indicate a statistically significant difference between BS OE and corresponding null lines as determined by independent t-test, where *p* ≤ 0.05 is flagged with one star (*), *p* ≤ 0.01 is flagged with 2 stars (**) and *p* ≤ 0.001 is flagged with three stars (***).
